# Exploring Meiosis in Brown Algae: Meiotic Axis Proteins in the model brown alga *Ectocarpus*

**DOI:** 10.1101/2024.12.18.629156

**Authors:** Emma I. Kane, Lioba S. Trefs, Lena Eckert, Susana M. Coelho, John R. Weir

**Affiliations:** Department of Algal Evolution and Development, Max Planck Institute for Biology, Tübingen 72076, Germany; Friedrich Miescher Laboratory of the Max Planck Society, Max-Planck-Ring 9, 72076 Tübingen, Germany

**Keywords:** meiosis, molecular evolution, brown algae, molecular modelling

## Abstract

Most extant eukaryotic systems share core meiosis-specific genes, suggesting meiosis evolved only once in the last eukaryotic common ancestor (LECA). These genes have been characterized as master regulators of meiotic recombination, ensuring genetically diverse lineages. However, our understanding is limited as eukaryotic organisms beyond the animal, plant, and yeast lineages remain poorly understood. Recently, core meiotic genes have been identified in the genome of the model brown alga *Ectocarpus*, but currently lack proper characterization. Here, we combine bioinformatic, structural, and biochemical approaches to characterise the axial element orthologs, meiotic *Ectocarpus* HORMA domain protein (ecHOP1) and its interactor, reductional division protein 1 (ecRED1), in order to elucidate the molecular mechanisms of synaptonemal complex (SC) and double strand break (DSB) formation in brown algae. We highlight novel domain architecture within ecHOP1 and ecRED1 that support HORMA domain conformational switches and quantify the thermodynamic parameters of these interactions. Together, our data suggests that brown algae may employ alternative HORMA domain regulation mechanisms compared with animal, plant and yeast systems, and provide clues for future studies on the evolutionary constraints and adaptation of meiosis across the tree of life.

## Introduction

Meiotic recombination is at the center of the staggering diversity of eukaryotic life on Earth (Bolcun-Filas & Handel 2018). Meiosis consists of two rounds of cell division, with no intervening S-phase. During meiosis I homologous chromosomes recombine giving rise to genetically distinct haploid gametes after meiosis II (Figure 1A) (Zickler & Kleckner 2023; Hunter 2015). Though core meiotic genes are generally conserved across eukaryotes (Thangavel *et al*. 2023; Arter & Keeney 2023; Chen & Weir 2024), their presence and functionality can vary among species. The presence of core meiotic genes suggests meiosis likely evolved only once from the last common eukaryotic ancestor (LECA) (Goodenough & Heitman 2014; O’Malley *et al*. 2019). Emerging model organisms have the potential to provide further insights into the conservation and diversity of meiotic processes. In this context, the brown algae, members of the Stramenopile lineage, have recently emerged as outstanding model organisms to study the universality or uniqueness of biological processes in a large evolutionary context (Coelho & Cock 2020). For instance, *Ectocarpus*, the model organism for brown algae (Coelho 2024), has a haploid-diploid sexual life cycle with dioicy (Figure 1B). These types of life cycle involve an alternation between haploid and diploid stages, with meiosis mediating the transition from diploid to haploid stages and syngamy reconstituting the diploid genome. Remarkably, these organisms can also reproduce via parthenogenesis of unfertilised gametes, leading to the development of a (haploid) organism that is capable of (apo)meiosis to transition to the gametophyte generation (Bothwell *et al*. 2010). Although the mechanisms underlying (apo)meiosis are currently unknown, this feature illustrates the plasticity of development in these fascinating yet underexplored organisms.

**Figure 1.**
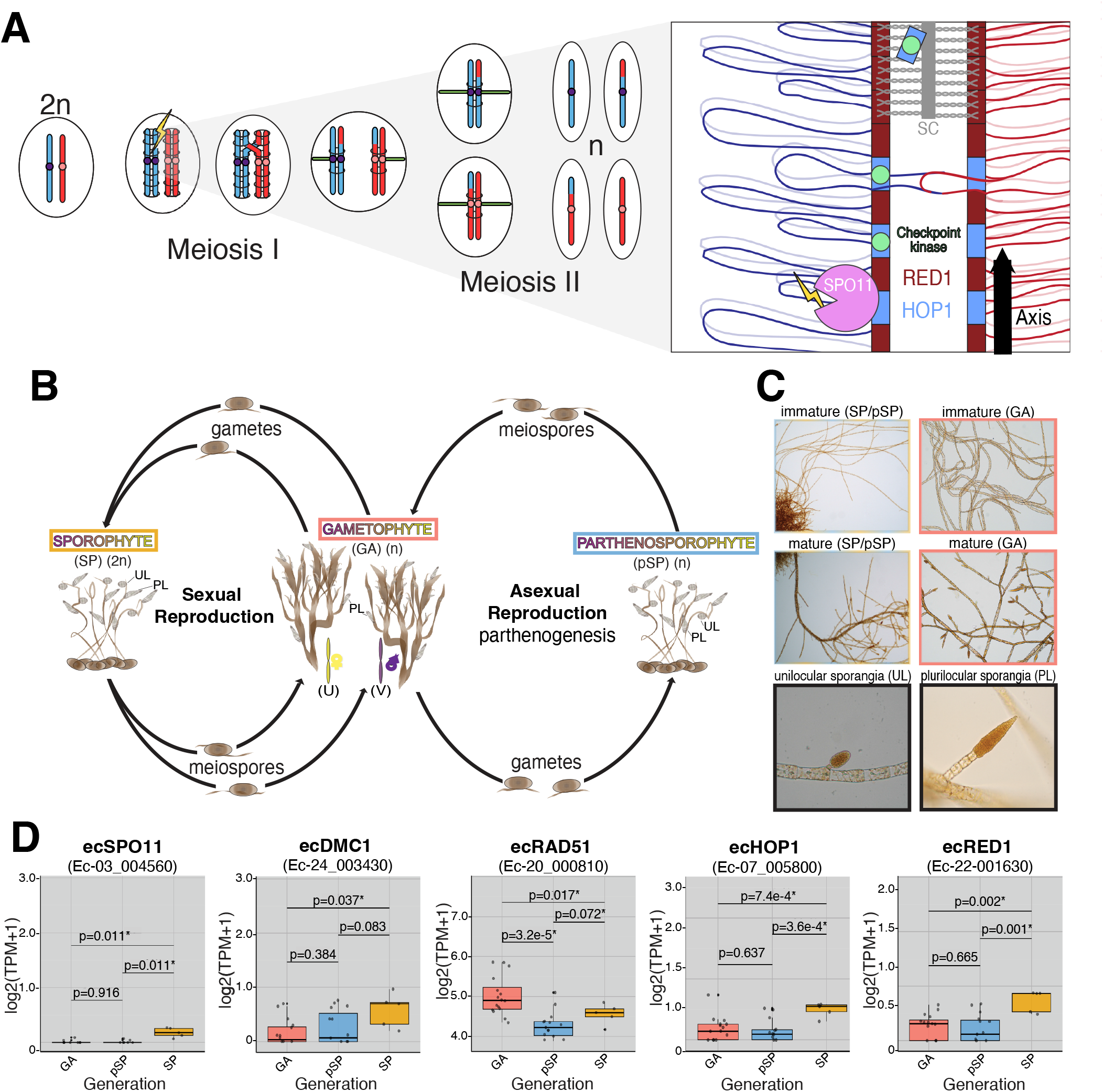
Meiosis, the *Ectocarpus* life cycle, and gene expression. A) Overview of Meiosis. A diploid cell (2n) with two homologous chromosomes (red and blue) undergoes DNA replication, where cohesin is loaded (black rings). Programmed double-stranded DNA breaks (yellow flash) are made, and repaired via homologous recombination to give rise to crossovers between homologs. At anaphase I arm cohesin is removed and homologous chromosomes segregated by the spindle (green). At meiosis II sister chromatids are separated after removal of pericentromeric cohesin. Inset Close up of meiotic axis between two homologous chromosomes. Red1 (deep red) recruits Hop1 (pale blue) to the axis. Hop1 in turn recruits Spo11 complexes that mediate DNA breaks (yellow flash). DNA damage activates the meiotic checkpoint via Hop1. Broken DNA strands are repaired from the homologous chromosome. Homologous chromosomes synapse, during which Hop1 is removed from the axis by Pch2 and the meiotic checkpoint is silenced. B) Overview of the *Ectocarpus* life cycle, which alternates between haploid (n) and diploid (2n) stages. Male and female gametes fuse to form a zygote, which develops into the sporophyte generation (SP; yellow). After meiosis in the unilocular sporangia (UL), the release of meiospores leads to the formation of gametophytes (GA; salmon), completing the sexual cycle. In the absence of male gametes, female gametes can undergo parthenogenesis, producing clones that develop into parthenosporophytes (pSP; cyan) which are morphologically indistinguishable from diploid sporophytes. These pSP produce (n) meiospores via a so far unknown process of (apo)meiosis (Bothwell *et al*. 2010). C) Microscopy images of immature and mature GA, SP, and pSP, alongside images of unilocular sporangia (unilocs, UL) and plurilocular sporangia (plurilocs, PL). The SP that successfully complete the sexual lifecycle to form GA are highlighted, with a focus on the PL and UL (where meiosis occurs). D) Boxplot showing the transcripts abundance of core meiotic and HORMA domain-containing genes, based on published RNAseq data. Data are presented for GA, pSP, and SP samples from male (Ec32) and female (Ec25) GA/pSP, as well as SP (Ec17) strains. Colors correspond to those in panels A and B, visualizing gene expression at different lifecycle stages. Statistical significance between samples is indicated (p < 0.05), and boxplots were generated in R.

In organisms studied to date, prophase I chromosomes have a distinct architecture consisting of regular loops of chromatin emanating from a proteinaceous axis (Grey & de Massy 2021; Wang *et al*. 2015; Kleckner 2006; Kleckner *et al*. 2003) (Figure 1A, inset). The meiotic axis is central to the control of recombination and crossover formation. The break-forming machinery assembles on the meiotic axis, whereas the breaks are made in the chromatin loops catalysed by the topoisomerase-like Spo11 (Blat *et al*. 2002). In yeasts and mammals the HORMA domain protein Hop1/HORMAD1 facilitates the connection between the DSB machinery and the axis (Stanzione *et al*. 2016; Dereli *et al*. 2024; Rousova *et al*. 2021). Hop1 is loaded onto the axis in a manner partially dependent on the AAA+ ATPase Pch2 (TRIP13 in mammals) (Raina & Vader 2020), where it binds to the axial component Red1 (SYCP2) (West *et al*. 2019). It is thought, though not formally proven, that Red1/SYCP2 is itself recruited to chromosomes via an interaction with cohesin complexes, due to both co-localisation on chromosomes, proteomics experiments, and the presence of cohesin binding motifs in Red1/SYCP2 (Sun *et al*. 2015; Köhler *et al*. 2017; Xu *et al*. 2019; Li *et al*. 2020). Hop1 has been shown to be phosphorylated in response to DSB formation by the ATM/ATR kinases (Carballo *et al*. 2008; Penedos *et al*. 2015). Phospho-Hop1 activates the meiotic checkpoint, which in yeast is mediated by the kinase Mek1, and controls progression through meiosis (Wu *et al*. 2010; Chen *et al*. 2018; Xu *et al*. 1997).

Pairing between homologous chromosomes manifests itself in the form of the synaptonemal complex, which forms along the length of the meiotic axis, possibly through a direct interaction between Zip1/SYCP1 and Red1/SYCP2 (Adams & Davies 2023). As chromosomes synapse, the meiotic axis is remodelled, suppressing further double-strand DNA break formation (Wojtasz *et al*. 2009). The length of the meiotic axis, relative to the size of the linear chromosome, has been shown to control the number of crossovers. Chromosomes with shorter axes have fewer crossovers, with longer axes more (Ruiz-Herrera *et al*. 2017). Therefore understanding the principles of meiotic axis assembly is key to understanding the initiation and regulation of meiotic recombination.

HORMA domain proteins play a critical role in DNA repair and checkpoint signalling (Gu *et al*. 2022). HORMA domains undergo conformational transformations from an ‘open’ to ‘closed’ state catalysed through interactions with closure motifs. Therefore, one HORMA domain can have distinct interactomes dependent on its topological state. Meiotic HORMA domain proteins (Hop1 in yeast, HORMAD1 in mammals, Asy1 in plants, HTP-1,2,3 and HIM3 in nematodes) have all so far been found to contain an N-terminal HORMA domain and one C-terminal closure motif, with the exception of HTP-3 which contains six closure motifs (Kim *et al*. 2014). Due to the presence of a cis closure motif it is expected that meiotic HORMA domains have a self-bound “ground-state”. Meiotic HORMA proteins can associate with the axis through a closure motif in other axial proteins (Red1 in yeast, SYCP2 in mammals, Asy3 in plants). Furthermore, a second recruitment pathway was recently shown for budding yeast Hop1, which can also associate directly with nucleosomes via a chromatin binding region (Milano *et al*. 2024; Heldrich *et al*. 2022). The chromatin binding region of Hop1 has been shown to be highly variable throughout evolution including complete absence in mammals and nematodes (Milano *et al*. 2024).

To better understand the conservation and diversity of meiotic protein function across eukaryotes, we investigated the role of meiotic proteins in *Ectocarpus. Ectocarpus* has been shown to have both a Spo11 ortholog (Brinkmeier *et al*. 2022), and brown algae possess a putative synaptonemal complex (Toth & Markey 1973). Moreover, candidate genes for both Red1 and Hop1 have been identified across a range of eukaryotes, including *Ectocarpus* (Tromer *et al*. 2021). Through genomic and structural analysis, we confirm the identity of these orthologs and biochemically characterise the core axial protein ecRED1 and its associated regulatory protein ecHOP1. We then characterise the DNA binding domain of ecHOP1 which binds preferentially to dsDNA, analogously to the nucleosome binding domain of budding yeast Hop1. Our findings reveal species-specific adaptations, including novel transcriptional isoforms and the presence of multiple closure motifs in ecHOP1. Together, our findings suggest that the meiotic axis in brown algae exhibits a blend of features seen in the meiotic axes of other eukaryotes, with functional plasticity arising from the unique utilization of a conserved set of meiotic genes.

## Results

### Lifecycle dependent expression of axial element orthologs

The life cycle of *Ectocarpus* involves an alternation between haploid male and female gametophytes (GAs) and diploid sporophytes (SP). Meiosis occurs in the unilocular (UL) reproductive structures in the sporophyte generation. *Ectocarpus* may also undergo an alternative asexual cycle via parthenogenesis, where female gametes develop autonomously (without fusion with a male gamete) into haploid parthenosporophytes (pSP). These pSP are morphologically indistinguishable from diploid sporophytes and at maturity undergo (apo)meiosis to produce haploid spores (meiospores). Therefore, in addition to a classical reductional meiosis, these organisms may undergo an alternative form of non-reductive meiosis (Bothwell *et al*. 2010) (Figure 1B and C).

We used published RNAseq data to investigate the transcript abundance of genes related to meiosis during the complex life cycle of *Ectocarpus* (Figure 1D). We focused on orthologs involved in recombination and synaptonemal complex (SC) formation. We found that SPO11, the conserved initiating factor for meiotic recombination (Bergerat *et al*. 1997; Keeney *et al*. 1997), was specifically upregulated in (diploid) sporophytes, where reductional meiosis occurs. Similarly, the meiotic-specific recombinase DMC1 (Bishop *et al*. 1992) showed a specific upregulation in diploid sporophytes. In contrast, RAD51, which is involved in both meiotic and somatic DNA repair, did not show an expression pattern specific to meiotic tissues (Figure 1D).

Previous bioinformatics work provided candidate genes for the axial protein RED1 (Tromer *et al*. 2021) (from hereon ecRED1) and its associated HORMA domain-containing partner HOP1 (ecHOP1). We therefore also investigated the transcript abundance of these genes. In line with the expression patterns for meiosis-specific SPO11 and DMC1, ecRED1 and ecHOP1 both showed specific expression in sporophyte tissue. ecHOP1 has two transcriptional isoforms reported through ORCAE (Ec_07-005800.1 and Ec_07-005800.1 respectively) (Sterck *et al*. 2012). Together, our data demonstrates that core meiotic genes are upregulated in stages of the life cycle where meiosis occurs, consistent with a role of these genes specifically in reductional meiosis but not in (apo)meiosis in the parthenosporophyte.

### Identification and computational analyses of ecHOP1

Both isoforms of ecHOP1 have a shared N-terminal region of the protein, while encoding a divergent C-terminal region (Figure 2A, Supplementary Figure 1A). We utilised isoform specific RT-PCR primers with RNA extracts from *Ectocarpus*, specifically unilocs collected during the SP lifestage and vegetative tissue (non-reproductive) from both SP and pSP lifestages. In these experiments we found direct evidence of expression of ecHOP1 isoforms 1 and 2 (Figure 2B).

**Figure 2.**
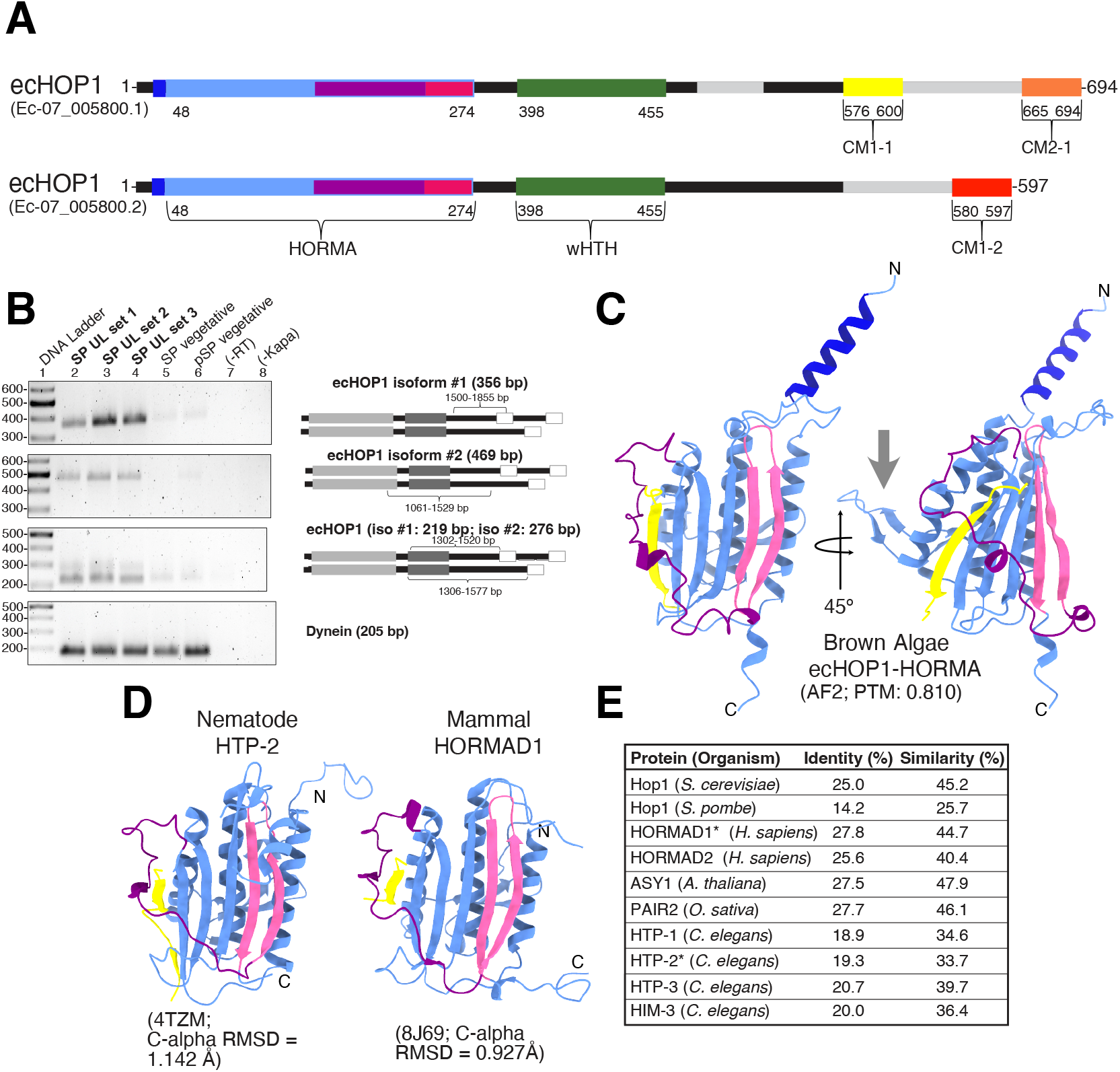
*Ectocarpus* HOP1 - Expression, Conservation and Structure. A) Domain architecture of ecHOP1 isoforms generated utilizing the sequences in ORCAE (Ec-07_005800.1 for isoform #1 and Ec-07_005800.2 for isoform #2), coupled with AlphaFold2 modeling. The boundaries for the putative CMs were determined from AlphaFold 2 models and alignment with previously identified CMs. Grey regions denote the variable sequences between the isoforms. Supplementary Figure **??**A shows the details of the differences between the isoforms. B) RT-PCR identified the presence of both isoforms from replicate uniloc samples collected from SP tissue (bold), and vegetative (non-reproductive) tissue from SP and pSP. Primer design was used to ensure specificity of amplified regions that are unique between both isoforms, as well as overlapping regions to identify both isoforms in the same sample (highlighted in the graphics below each result). RT-PCR controls, as well as amplifying for the housekeeper gene Dynein were utilized. C) AlphaFold2 model of the EcHop1 HORMA domain (light blue) bound in cis to a predicted closure motif (yellow). The N-helix is shown in dark blue, the safety belt in purple and the buckle in pink. The β3 hairpin is denoted with a grey arrow. (D) Experimentally determined HORMA domain structures from nematodes (*C. elegans*; HTP-3 with HIM-2 CM (PDB: 4TZM)) and mammalian (H. sapiens HORMAD1 bound in cis to HORMAD1 CM (PDB: 8J69)) PTM. E) Sequence identity and similarity determined from pairwise alignment of the *Ectocarpus* HORMA domain with the meiotic HORMA domains from various model organisms. Pairwise alignments were evaluated using the EMBOSS Needle Job Dispatcher through EMBL-EBI(Madeira *et al*. 2024).

In the absence of experimental structural information on ecHOP1 we utilized AlphaFold2 (Jumper *et al*. 2021) to gain insights into the structure and function of the ecHOP1 isoforms. Both isoforms are predicted to contain a conserved N-terminal HORMA domain and a winged helix-turn-helix (wHTH) domain, which may bind to chromatin, as in budding yeast Hop1 (Milano *et al*. 2024) (Figure 2A, Supplementary Figure 1B). The predicted HORMA domain of ecHOP1 (aa. 50-274) has structural similarity with meiotic HORMA domains from nematode (PDB 4TZM (Kim *et al*. 2014)) and mammal (PDB 8J69 (Wang *et al*. 2023)) orthologs (Figure 2C), with a C-alpha RMSD of 1.142 Å and 0.927 Å respectively (Figure 2D). ecHOP1 also has a high level of sequence similarity with other identified meiotic HORMADs (Figure 2E).

Close inspection of the AF2 structure of the ecHOP1 HORMA domain revealed two features that have not previously been described in other meiotic HORMA domains. Firstly, we observe an extended alpha-helix in the N-terminal region, which we term N-helix (Figure 2C, blue). We also find a beta-hairpin between residues 94 and 103, inserted between the β3 and αB regions (Figure 2C grey arrow, Supplementary Figure 1B and C). We term this region the β3”hairpin, which is conserved within brown algae (Supplementary Figure 1C).

### Biochemical characterization of ecHOP1-HORMA

To investigate ecHOP1 further we produced recombinant ecHOP1 HORMA domain. Given our observation that ecHOP1 also contains an N-terminal extended alpha-helix (N-helix, Figure 2C) we utilized a construct with the full N-terminal region consisting of residues 1-274, which we call ecHOP1-HORMA. We expressed ecHOP1-HORMA in E. coli with an N-terminal 2xStrep-II and purified the protein to homogeneity (Figure 3A, Supplementary Figure 2A). Size-exclusion chromatography coupled with multiangle light scattering (SEC-MALS) revealed a species of ecHOP1HORMA at 34.01 kDa (+/-7.5%) (Figure 3B), consistent with a monomeric ecHOP1HORMA (theoretical Mw of 36.19 kDa). We also observed a second species, with mass of 57.52 kDa (+/-31.3%), which could represent a small dimer fraction. Other HORMA domain proteins, for example Mad2 and Rev7 are known to form homo- and heterodimers via their HORMA domains (Supplementary Figure 3A), though this has never been observed for a meiotic HORMAD.

**Figure 3.**
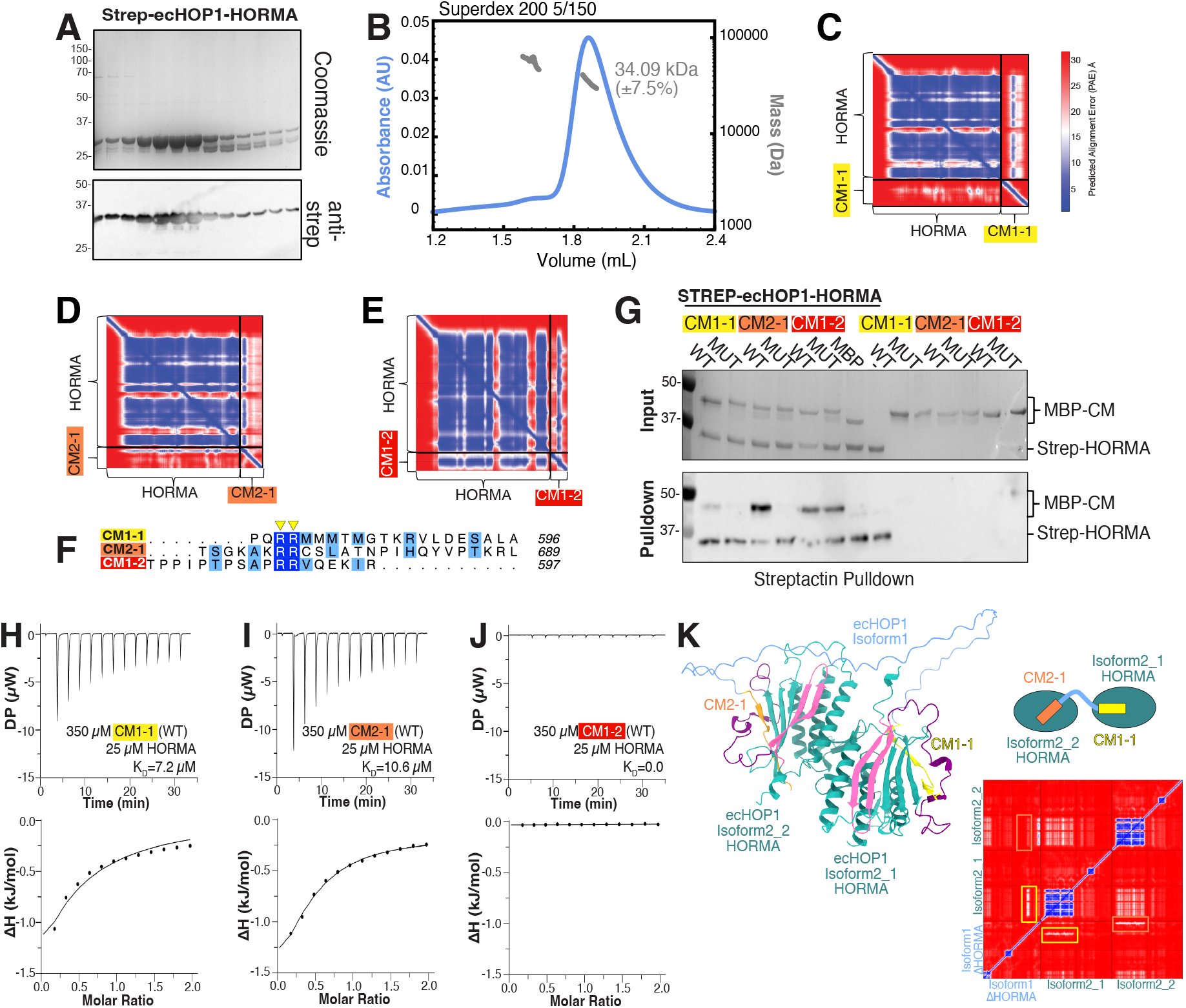
in-vitro characterization ecHOP1-HORMA with HOP1 closure motifss. (A) Upper panel - coomassie stained SDS-PAGE gel of a final size exclusion purification of ecHOP1-HORMA. Lower panel - anti-Strep-II western blot of the same fractions. The corresponding chromatogram can be found in Supplementary Figure 2A (B) SEC-MALS of ecHOP1-HORMA run on a Superdex 200 5/150 column. Blue trace shows the absorbance at 280 nm in arbitrary units (AU). The gray trace shows the molecular mass measurement. The main peak corresponds to a measured MW of 34.09 kDa. The second peak has a mass of 57.5 kDa. (C-E) Predicted alignment error (PAE) plots of AlphaFold2 predictions of ecHOP1-HORMA with CM1-1, CM2-1 and CM1-2. F) Aligned ecHOP1 closure motifs with the conserved arginine residues highlighted with yellow triangles. G) Pull-Down assays with the WT and mutant (R/A) CMs of ecHOP1. 2xStrepII-ecHOP1-HORMA was used as bait for the prey MBP-tagged CMs. In addition MBP-CMs were used alone as a control for background binding to the Streptactin beads. H-J) Isothermal titration calorimetry (ITC). 350 µM of indicated MBP-tagged closure motif was titrated against 25 µM of ecHOP1-HORMA in the cell. Buffer-buffer controls were run and subtracted from the experimental data to yield the heats shown. Binding curves were fitted in the software and the determined *K*_*D*_ is shown. K) AlphaFold2 model of a complex of ecHOP1-isoform1 (with the N-terminal HORMA domain removed), and two copies of ecHOP1-isoform2. The model is coloured as elsewhere, but the isoform2 HORMA domains are coloured in teal. In the PAE plot the CMs are highlighted with the colours as in the cartoon.

AlphaFold2 modeling of an ecHOP1 HORMA dimer (Supplementary Figure 3B), indicated moderate confidence for dimer formation. Closer analysis of predicted dimer structures suggested homodimerization of ecHOP1 may be limited by steric hindrance from the novel N-helix (Supplementary Figure 3C). Indeed upon removal of N-helix, the confidence in the dimer model increased (Supplementary Figure 3D). We carried out pull-down assays using N-terminal tagged HORMA domains (StrepII and His6SUMO) to prevent crossreactivity during Western blot analysis and assay for potential self-association. We also carried out pull-downs using ecHOP1HORMA lacking the N-helix (ecHOP150-274), nonetheless we observed no definitive evidence of HORMA-HORMA dimerization for ecHOP1-HORMA (Supplementary Figure 3E).

### Interaction of ecHOP1-HORMA with ecHOP1 CMs

Since previously studied meiotic HORMADs interact with their own CMs - located in the extreme C-termini of meiotic HORMADs - in cis, we asked whether AlphaFold2 could predict potential CMs in both of the ecHOP1 isoforms. Curiously, ecHOP1 isoform 1 was predicted to contain two potential CMs (CM1-1 and CM2-1 comprising residues 567-600 and 665-694 respectively), and isoform 2 (CM1-2) in the very C-terminal region (residues 580-597) (Figure 3C-E) These CMs contain conserved arginine residues, potentially important for HORMA domain interaction as has previously been seen in other CMs (West *et al*. 2017; Kim *et al*. 2014) (Figure 3F) but are not otherwise very similar to one another. This sequence degeneracy has previously been observed for other meiotic closure motifs (West *et al*. 2017; Kim *et al*. 2014).

To test potential interaction between ecHOP1-HORMA and the three predicted CMs (Figure 2A, Figure 3F), we produced recombinant ecHOP1 CMs fused to an N-terminal maltose binding protein (MBP) to improve solubility. For each CM we produced a mutant variant where the arginine residues were mutated to alanine (Figure 3F). We first tested the interaction between Strep-II tagged ecHOP1-HORMA and MBP-CMs in a pull-down on Strep Tactin-XT beads (Figure 2A & Figure 3G). Here we observed an interaction for all three CMs. For CM1-1 and CM2-1 the binding was also disrupted by the introduced mutations, but not for CM1-2, where both the wild-type and mutant sequences appeared to bind to ecHOP1-HORMA.

To gain further insights into CM to ecHOP1HORMA interactions we determined the relative binding affinities by isothermal titration calorimetry (ITC) (Figure 3H-J). Under these conditions we could further confirm the interaction of CM1-1 and CM2-1 with ecHOP1HORMA. The measured dissociation constants (*K*_*D*_) for CM1-1 and CM2-1 were 7.2 and 10.6 µM respectively. Curiously we did not detect an interaction for CM1-2. Based on this, and that the mutations in CM1-2 did not disrupt binding in the pull-down experiment, we conclude that CM1-2 is not a functional CM, and that ecHOP1 isoform 2 does not contain a CM for cis-binding. This raises the possibility that ecHOP1-isoform1 can recruit two ecHOP1-HORMAs. To test this idea further we utilised AlphaFold2 to model ecHOP1 isoform 1 (without the N-terminal HORMA domain) together with two copies of ecHOP1 isoform 2 (Figure 3K). This model shows CM1-1 and CM2-1 bound in cis to two copies of ecHOP1-isoform 2 (Figure 3K).

### DNA binding capabilities of ecHOP1

As described above AlphaFold modelling of ecHOP1 predicted the presence of a winged helix-turn-helix (wHTH) domain (a.a. 398-455). This domain is conserved among HOP1 in Stramenopiles (Supplementary Figure 4A), though occasionally as a twin wHTH domain (Milano *et al*. 2024). An expanded version of the wHTH domain in budding yeast (which also contains a PHD domain) was recently found to bind to nucleosomes (Milano *et al*. 2024). To investigate the chromatin binding properties of the wHTH domain we produced recombinant SUMO-tagged ecHOP1-wHTH, and purified this to homogeneity (Supplementary Figure 2B) and evaluated DNA binding via EMSAs.

We first asked whether ecHOP1-wHTH might be able to bind to mononucleosomes wrapped in 167 bp Widom DNA (Lowary & Widom 1998), as has been shown for the equivalent region of *S. cerevisiae* Hop1 (Milano *et al*. 2024). However, we only observed weak binding under these conditions (Figure 4A). We then tested whether wHTH could bind to the 167 bp Widom dsDNA alone and observed robust binding, for which we determined an apparent *K*_*D*_∼250 nM (Figure 4B and C). As a comparison we also measured DNA binding for ecHOP1HORMA, which did not show any clear DNA binding (Figure 4D).

**Figure 4.**
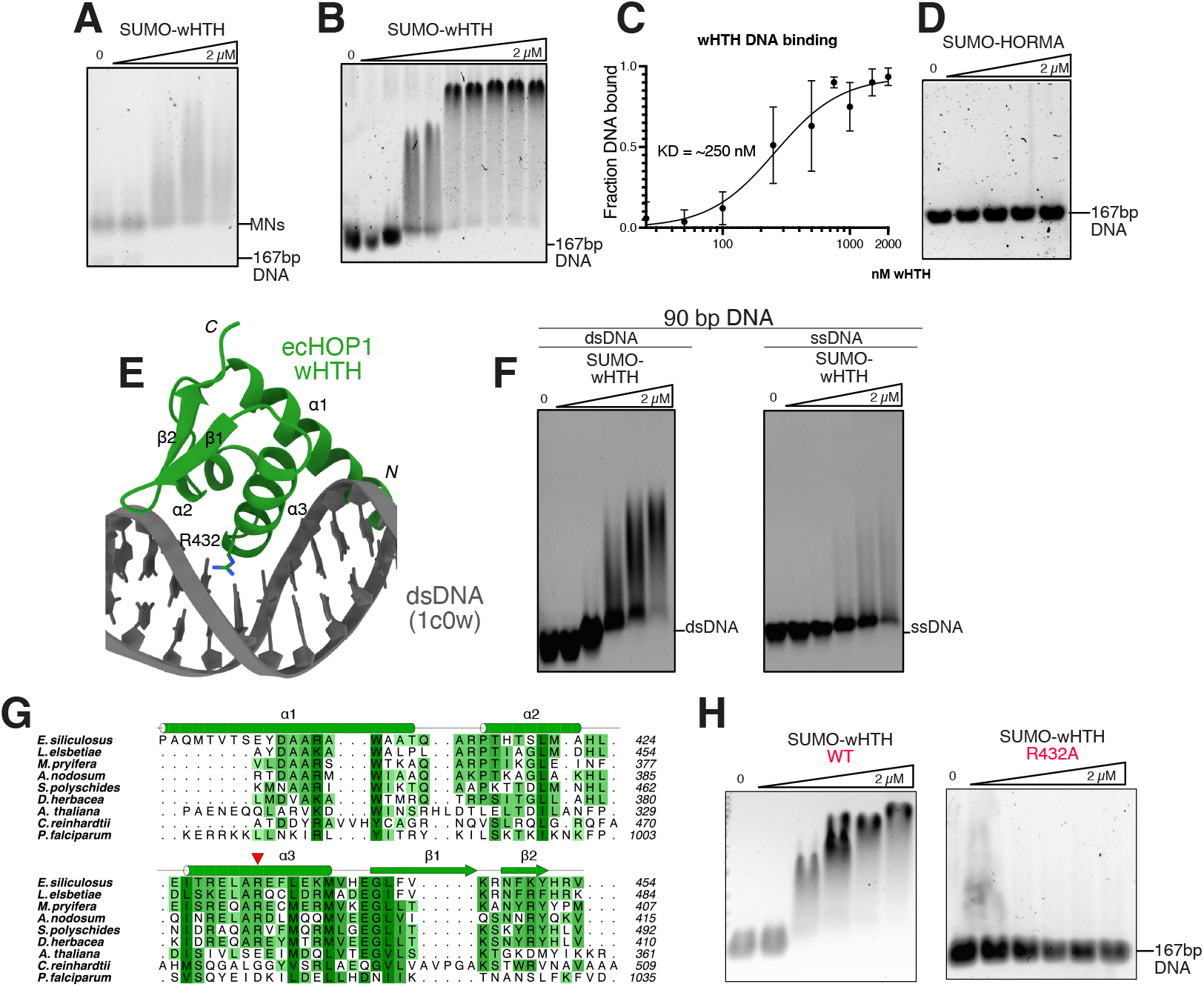
DNA binding properties of the ecHOP1-wHTH domain. A) EMSA of SUMO-tagged ecHOP1 wHTH titrated against 50 nM of nucleosomes wrapped in 167 bp Widom DNA. B) EMSA of SUMO-tagged ecHOP1 wHTH titrated against 50 nM of 167 bp DNA. C) Binding curve based on triplicate experiments as shown in B). Curve fitting carried out in Prism required the use of the Hill Coefficient for the best fit (h=1.7) negative control EMSA with the cleaved His6-SUMO fusion tag, to demonstrate if the tag influences DNA binding. Here no binding can be observed. D) ecHOP1-HORMA domain was also titrated against 167 bp Widom DNA in an EMSA experiment as a negative control. E) Composite structural model. The AlphaFold2 prediction of ecHOP1-wHTH (residues 398-454) superimposed onto the wHTH in PDB ID 1C0W. The dsDNA from 1C0W is shown (grey) and the model should approximate the DNA binding mode of ecHOP1-wHTH. R432 of ecHOP1 sits in the middle of the candidate DNA binding helix, and is proximal to the sugar-phosphate backbone of the dsDNA. F) EMSAs of ecHOP1-SUMO-wHTH binding to 90-mer dsDNA and ssDNA. G) MSA of multiple wHTH domains from across Stramenopiles in addition to *A. thaliana, C. reinhardtii* and *P. falciparum* sequences. The location of R432 of ecHOP1 is indicated by the red arrow in the middle of the α3-helix. H) EMSA showing the titration of ecHOP1-SUMO-wHTH R432A mutant against 167 bp Widom DNA.

We carried out a DALI search with the AlphaFold2 model of ecHOP1-wHTH to find structurally similar wHTH domains in complex with nucleic acids in the PDB. We focused on the structure of DtxR with dsDNA (PDB ID 1C0W). We superimposed the AF2 model of ecHOP1-wHTH onto DtxR with dsDNA, which gave a C-alpha RMSD of 0.727 Å (Figure 4E). In this structure the wHTH binds to dsDNA via the major groove, with α3 of wHTH sitting in the major groove. If ecHOP1-wHTH is binding via a similar mechanism we reasoned that this would strongly prefer dsDNA over ssDNA. We evaluated this in an EMSA using a 90-mer ssDNA and dsDNA, and found that ecHOP1-wHTH indeed binds more tightly to dsDNA (Figure 4F).

We produced a MSA of HOP1-wHTH from Stramenopiles, and added previously identified Hop1 wHTH domains from A. thaliana, C. reinhardtii and P. falciparum (Figure 4G, Supplementary Figure 4A). By combining the AF2 model superimposed onto dsDNA and the MSA we identified residues within α3 of the ecHOP1-wHTH domain that could mediate DNA binding. R432 is positioned within α3 the helix, and is proximal to the sugar-phosphate backbone of the DNA. We produced mutant recombinant ecHOP1-wHTH with an R432A mutation. In an EMSA experiment we detect no dsDNA binding with ecHOP1-wHTHR432A when compared with (Figure 4H, Supplementary Figure 4 B and C). These results establish ecHOP1-wHTH binds preferentially and robustly to dsDNA, likely via interactions mediated by α3, with R432 playing a critical role in this binding, as demonstrated by the loss of DNA interaction in the R432A mutant.

### Characterization of the ecHOP1 interactor ecRED1

Next, we expanded our studies to include the predicted Red1 ortholog in *Ectocarpus*, (from here on ecRED1). Analysis of the AlphaFold2 predicted structure of ecRED1 shows that the protein contains an N-terminal armadillo (ARM) domain, a mid-region Pleckstrin homology (PH) domain, and a C-terminal coiled-coil (CC) domain (Figure 5A, Supplementary Figure 5A), a domain arrangement widely found conserved in predicted Red1 orthologs (Feng *et al*. 2017; West *et al*. 2019). The coiled-coil domain of Red1 (yeast) and SYCP2 (human) has been shown to form homotetramers (West *et al*. 2019). AlphaFold2 modelling of the ecRED1 coiled-coil region suggests that this could also be the case for ecRED1 (Supplementary Figure 5B).

**Figure 5.**
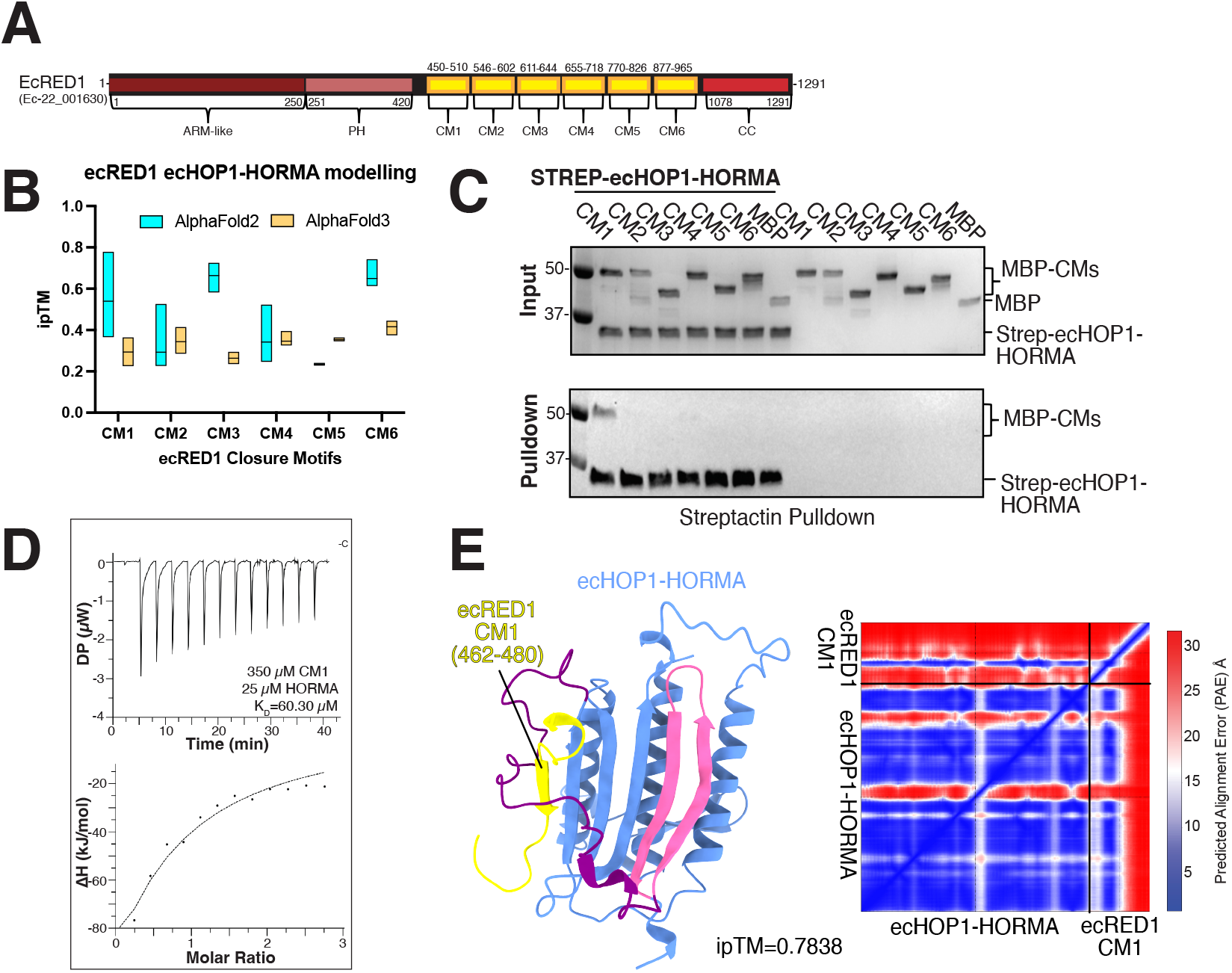
Characterization of ecRED1 closure motif interactions with ecHOP1-HORMA. (A) Domain architecture of ecRED1, with the sequence from ORCAE (Ec-22_001630), coupled with domain boundaries determined via AlphaFold2. The ARM-like, PH and CC domains were modelled individually in AlphaFold2 (Supplementary Figure 5A). Determination of multiple CMs in ecRED1 was made via MSA and AlphaFold2 modeling, where multiple models were generated with truncations of ecRED1. B) Summary of ipTM values (interchain confidence score) based on the top 5 scores from 25 predictions for both AlphaFold2.3.2 and AlphaFold3 (run on www.alphafoldserver.com) The line in the bar plot shows the median value for the predictions (C) Pull-Down assays with 2xStrepII-ecHOP1-HORMA domain and the putative MBP-tagged CMs on Streptactin beads. (D) 350 µM of MBP-CM1 was titrated against 25 µM ecHOP1-HORMA in the cell. A buffer-buffer control experiment was subtracted from the data to provide the heats shown. A *K*_*D*_ of 60.3 µM was determined. E) AlphaFold2 model of ecHOP1-HORMA in complex with ecRED1-CM1 (yellow). The predicted alignment error plot for the model is also shown.

In other species, Red1 orthologs have been shown to contain a CM that binds to the HORMA domain of Hop1 ortholog (West *et al*. 2019; Woltering *et al*. 2000). To find potential CMs in ecRED1 we carried out AlphaFold2-multimer (Evans *et al*. 2021) predictions of ecHOP1-ecRED1 sequences. The algorithm predicted six potential CMs, with varying levels of confidence based on the interchain confidence score (ipTM) (Figure 5B). We also ran our candidate closure motifs using AlphaFold3 (Abramson *et al*. 2024). Here the algorithm gave overall lower ipTM scores, and with no differences between the CMs (Figure 5D). We therefore undertook a biochemical approach to validate the ecRED1 CMs. Curiously the number of ecRED1 CMs, six, is also the same number of CMs found in *C. elegans* HTP-3 (Kim *et al*. 2014).

We expressed and purified all predicted ecRED1 CMs as N-terminal MBP-fusion proteins. In pull-downs on StrepII-tagged ecHOP1, ecRED1 CM1 demonstrated binding to the ecHOP1 HORMA domain, and although faint, CM4 was also observed as a putative interactor (Figure 5C). We then performed a phylogenetic analysis of the ecHOP1 and EcRED1 CMs across the brown algal family, which revealed that CM motifs such as “(K/R)MMMT(K/R)” in CM1-1 and “(K/R)CSLA(K/R)” in CM2-1 are highly conserved (Supplementary Figure 6).

ITC results showed that CM1 exhibited a binding affinity of 60 µM (Figure 5D). This is nearly 10x weaker than CM1-1 of ecHOP1 (Figure 3H). Curiously, this is inverse to observations from yeast, where RED1 CMs exhibit stronger binding to HORMA domains than HOP1 CMs (West *et al*. 2017). The remaining putative ecRED1 CMs could not be validated through *in vitro* analyses. The high conservation of the ecRED1-CMs within the brown algal family renders the possibility that under specific biological conditions they may either act as CMs, or have another important function (Supplementary Figure 6).

Taken together, our findings reveal novel insights into brown algal meiotic regulation, emphasizing the intricate interactions between HORMA domains and CMs in proteins like ecHOP1 and ecRED1. A proposed model for meiotic regulation in brown algae, based on these interactions, is presented in Figure 6.

**Figure 6.**
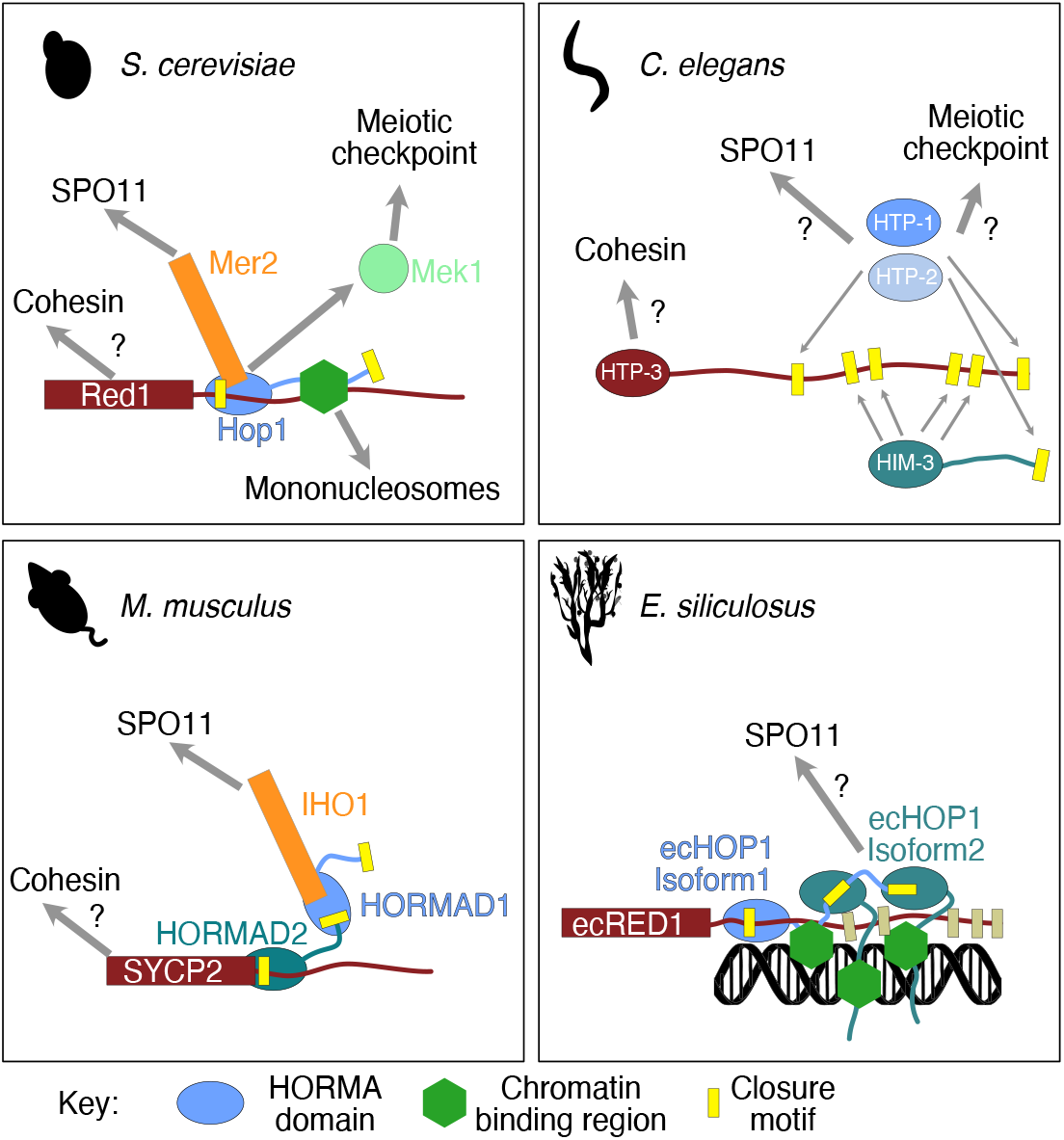
Comparison of the *Ectocarpus* axial proteins with other model systems. Red1 contains one CM, which can bind to Hop1. Hop1 can be recruited partly independently of Red1 via its CBR binding to nucleosomes(Milano *et al*. 2024; Heldrich *et al*. 2022). Hop1 can directly bind to Mer2 when it is bound to the CM of Red1(Rousova *et al*. 2021), which in turn regulates Spo11 recruitment. When Hop1 is phosphorylated it can bind to the checkpoint kinase Mek1 (Carballo *et al*. 2008; Hollingsworth & Ponte 1997). In *C. elegans* there is no Red1 ortholog, instead HTP-3 is recruited to the axis, possibly via cohesin interaction where it can bind to both HIM-3 and HTP-1 and HTP-2 via six closure motifs(Kim *et al*. 2014). In mammals the SCYP2 protein can bind directly to HORMAD2 via a closure motif(West *et al*. 2019). SYCP2 might also bind to cohesin, via the CTCF binding site (Li *et al*. 2020). The CM of HORMAD2 can bind to HORMA1, which in turn can bind to the phosphorylated C-terminus of IHO1 (Stanzione *et al*. 2016; Dereli *et al*. 2024). IHO1 is then able to assemble further SPO11 activating factors, and recruit/activate SPO11 (Dereli *et al*. 2024; Laroussi *et al*. 2023). In E. siliculosus, ecRED1 can also bind to ecHOP1-HORMA via CM1. ecRED1 also contains several putative CMs (greyed out boxes), which may be activated via post-translational modifications. Here we show ecHOP1-isoform1 recruiting 2 x ecHOP1-isoform2 proteins via CM1-1 and CM2-1. Alternative configurations of ecHOP1 are presumably also possible, which may lead to larger branched assemblies of ecHOP1 on chromatin. In addition to recruitment via ecRED1, ecHOP1 also contains a chromatin binding region in the form of a dsDNA binding wHTH domain (green hexagons).

## Discussion

Meiosis is a ubiquitous feature of eukaryotic life, yet even among conventional model organisms, there is considerable diversification of the molecular machinery underpinning meiosis (Arter and Keeney 2023). Utilizing tissue specific transcriptomic data, we have confirmed the meiotic expression of two chromosomal associated proteins ecHOP1 and ecRED1. Structural modeling combined with biochemical and biophysical assays suggest that these proteins share similarities with orthologs in other eukaryotic organisms, but also reveal unique features.

Given that, in other species, Hop1 is a regulatory factor for the formation of meiotic double-stranded DNA breaks (DSBs) we wanted to understand how ecHOP1 might be localized to chromosomes. We find two possible pathways. One is via a direct interaction between the closure motif in ecRED1, a second is via the DNA binding properties of the wHTH domain (Figure 4, Figure 6). Unlike in budding yeast Hop1, we observe no nucleosome binding for ecHOP1 (Figure 4A). Instead, we find that ecHOP1-wHTH binds to double-stranded DNA. While AlphaFold2 modelling suggested ecRED1 might contain six CMs, we could only biochemically validate one of these (Figure 5 C,D). Nonetheless, Red1 in other species (for example, in budding yeast) is highly post translationally modified (Kar *et al*. 2022). If this is also the case in brown algae, these additional putative CMs could be functional under certain circumstances, raising the level of ecHOP1 recruitment at certain loci (Figure 6).

AlphaFold2 modelling suggested three possible ecHOP1 cis-CMs, two in isoform 1 (CM1-1 and CM2-1) and one in isoform 2 (CM1-2). While AlphaFold2 predicted CM1-2 with the highest confidence, we found that both CM1-1 and CM2-1 bind to the HORMA domain with equivalent affinity (Figure 3E). Bioinformatics analysis shows that CM1-1, CM2-1, and ecRED1 CM1 are common to other brown algal lineages (Supplementary Figure 6). ecHOP1 also expresses a second isoform, which contains a HORMA domain, a DNA binding domain, but no closure motif. Taken together this suggests a possible arrangement of ecHOP1-isoform1 being recruited to ecRED1-CM, whereupon it exposes two functional CMs, which could then bind to further HORMA domains (Figure 3K, Figure 6). These further HORMA domains could be ecHOP1-isoform1 or isoform2, leading to different possible branched structures. Likewise, isoform2 could be recruited to ecRED1, which would not recruit any further ecHOP1. While the “beads on a string” model for meiotic HORMA domain recruitment has been previously proposed, this type of complex branching assembly of meiotic HORMA domains has previously only been proposed in Nematodes, which contains multiple meiotic HORMA domains. These potential arrangements, combined with the ecRED1 cryptic closure motifs suggest that *Ectocarpus* might utilise varying levels of chromosomal ecHOP1 as a means of regulating DSB formation and the meiotic checkpoint.

This study also highlights the need for biochemical validation of computational models. In the case of the HORMA domain to CM interactions, all putative CMs were predicted to bind to the HORMA with similar confidence by both AlphaFold2, and by AlphaFold3 (although the latter provided consistently lower confidence scores). Biochemical assays could only validate 3 of the 9 predicted CMs, and one of these required a quantitative assay to disprove binding to the closure motif. It remains to be seen if these predicted CMs are spurious, or if they are activated through post-translational modifications.

Our work raises a number of further questions. Firstly, how is ecRED1 recruited to the axis? Given that ecRED1 appears to be similar to *S. cerevisiae* Red1 and mammalian SYCP2 we might predict that it would bind to cohesin, possibly in a manner similar to binding by CTCF (Li *et al*. 2020). Secondly, how would ecHOP1-isoform1 be recruited to ecRED1 if it is bound to its own CM? Again, we might speculate that a Pch2/TRIP13 AAA+ ATPase plays a role in opening the HORMA domain and displacing the CM (Raina & Vader 2020). Thirdly, what are the downstream interactors of ecHOP1? Given the presence of ecSPO11 (Brinkmeier *et al*. 2022), we speculate that there are additional factors, similar in function to IHO1/Mer2 that help to recruit and activate SPO11. Furthermore, since we note that ecHOP1-HORMA has additional structural elements; the N-helix and the β3 hairpin, it might utilise these to bind to a wider range of downstream factors than found in other species. Since the AAA+ ATPase Pch2/TRIP13 interacts with the Hop1 HORMA domain via the N-terminus, it is tempting to speculate that the N-helix may regulate this process, and further work will be required to test this idea. Finally, what is the identity of the checkpoint kinase in *Ectocarpus*? So far, only Mek1 has been unambiguously assigned as the meiotic checkpoint kinase in budding yeast. Further work remains to determine if *Ectocarpus* also utilises a meiosis specific checkpoint kinase, or another cellular kinase fulfils this function.

By tracing the evolutionary lineage of these proteins, our study provides a framework for future research into the molecular evolution of meiosis. It highlights the potential for uncovering regulatory mechanisms in other non-major model organisms that could offer further insights into the origins and diversification of meiosis. In this way, our work not only deepens our understanding of meiotic recombination but also opens new avenues for exploring how evolutionary pressures have shaped the molecular machinery of life over large timescales.

## Materials and Methods

### RNAseq and Statistical Analysis

Published RNAseq datasets (Lipinska *et al*. 2017; Lotharukpong *et al*. 2024) were processed to visualize the expression levels of the core meiotic genes across the *Ectocarpus* life stages (GA, SP, and pSP). Reads from the diploid Ec17 strain were used for SP and Ec32 and Ec25 (haploid strains) were used for GA and PSP stages. Raw counts were transformed into log-transformed expression values as log2(TPM + 1), where TPM stands for Transcripts Per Million. Boxplots were generated to display the distribution of log-transformed expression levels across three tissue groups: GA (gametophyte), pSP (parthenosporophyte), and SP (sporophyte). To ensure consistency, specific colors were assigned to each group: salmon/magenta for GA, blue/cyan for pSP, and mustard yellow for SP. The boxplots were generated using the ggplot2 package in R. Each gene was visualized individually, with jitter plots added to display individual data points. Custom y-axis limits were calculated for each gene, and plots were presented with a grey background and muted axis/grid lines for visual clarity. Pairwise comparisons between the three tissue groups (GA vs. pSP, pSP vs. SP, and GA vs. SP) were performed using two-sided t-tests. An overall analysis of variance (ANOVA) test was conducted to determine if there were significant differences among the three groups for each gene. P-values were annotated on the boxplots, and statistical significance was assessed between groups. Brackets were manually added to indicate significant differences between tissue groups, with the corresponding p-values displayed adjacent to the brackets. All statistical analyses were conducted using R version 4.3 with a significance threshold set at p < 0.05.

### RNA extraction from brown algal tissue

Freshly cultivated *Ectocarpus* strains were prepared as previously described (Coelho *et al*. 2012). Algae were flash frozen in liquid nitrogen placed into individual 1.5 mL Eppendorf tubes, and homogenized into a fine powder with tissue grinding pestles (BioEcho Life Sciences). Throughout the lysis, the tubes were routinely placed back into LN. 50 µL of preheated CTAB3 Extraction Buffer (100 mM Tris-HCl, pH 8.0, 1.4 M NaCl, 20 mM EDTA, 2% plant RNA isolation aid (PVP), 2% CTAB, 1% β-mercaptoethanol) was added to each sample and vortexed for one minute, and quickly centrifuged for a one second pulse. The lysed solution was then transferred to a RNAse-free 2 mL tube (Eppendorf). 50 µL of achloroform/isoamylalcohol (24:1) solution was added, vortex vigorously, and then subjected to centrifugation at 10000 RPM for 15 minutes at 4 °C, following the segregation of the upper aqueous into a new RNAse-free 2 mL tube. This process was repeated twice. On the third cycle, the upper aqueous phase was transferred into a new 1.5 mL RNAsefree tube (Eppendorf) with 100 µL of 7.5 M LiCl and a 1% (v/v) β-mercaptoethanol. The samples were agitated vigorously and RNA precipitated O/N at -20 °C. The samples were centrifuged at 15K rpm for one minute at 4 °C, and the supernatant was discarded. The pellet was washed with 1 mL of ice cold 70% ethanol, then centrifuged again at 5K rpm for 10 minutes at 4 °C. The residual ethanol was removed from the pellet, and the samples were left to dry for no more than five minutes. The pellets were dissolved in RNAse-free H2O and RNA concentration was determined with a Nanodrop spectrophotometer. DNAse treatment was completed with the TURBO DNAse Kit (ThermoFisher) following the manufacturer’s protocol. RNA aliquots at 500 ng were prepared and stored in -80 °C for future use.

### RT-PCR to validate ecHOP1 transcriptional isoforms

Tissue samples (in triplicate) of Ec17 (SP lifestage) unilocular sporangia (uniloc sets 1-3), SP vegetative tissue (without unilocular sporangia), and Ec32 (pSP) vegetative tissue (without unilocular sporangia) were utilized for RT-PCR to determine RNA expression of both ecHOP1 isoforms. RNA extraction followed a protocol, previously optimized for *Ectocarpus* tissue (ref. 10.1038/s41559-022-01692-4). Complementary primers were designed with consideration for the difference in exons between each isoform, as well as control ecHOP1 primers that are specific to regions that encapsulate 100% conservation between both isoforms (Supplementary Table 1). Dynein was utilized as a control since it is a house-keeping gene. cDNA lacking reverse transcriptase (-RT) and Kapa polymerase (-Kapa) of pSP vegetative tissue were also used for PCR negative controls. Reactions were run under the “Touchdown” protocol to increase primer specificity. Following amplification, 60 DNA Loading Dye (ThermoFisher) was added into each sample at a 1:5 ratio and loaded onto a 2% agarose gel. The gel ran at 80 V for approximately one hour.

### AlphaFold modeling

Modeling of protein structures was performed with the AlphaFold multimer system (version 2.3.2) (Jumper *et al*. 2021; Evans *et al*. 2021) with two multimer predictions per model and the Amber relaxation procedure applied to the best model (Case *et al*. 2023). While modeling the ecHOP1 HORMA domain together with ecRED1 to discover the ecRED1 CM, several putative ecRED1 CMs appeared in the predicted models. First, the HORMA domain of ecHOP1 was modeled together with full-length ecRED1, followed by the model generations of the HORMA domain with ecRED, however with a truncation of the C-terminal CC region to mitigate low confidence model predictions due to limitations in CC domain predictions. After evaluating all putative CMs consistently appeared within the same boundaries in these models as well, further predictions were generated solely with the truncations of each putative CM in conjunction with the HORMA domain.

### CM Motif Identification and analysis

The phylogenetic tree used to compare the evolution of HOP1 and RED1 CMs in brown algae was modelled after a reference previously reported. Protein sequences of different brown algae species were retrieved through the BLAST interface of the Phaeoexplorer brown algal genome database (https://phaeoexplorer.sb-roscoff.fr/home/), using the brown algal protein sequences of ecHOP1 and ecRED1 as query. Results of the BLAST search were aligned using MAFFT MSA tools(Katoh *et al*. 2002). Manual curation of the MSA was performed using JalView (Waterhouse *et al*. 2009), and alignments for each putative CM were determined by N- and C-terminal K and/or R residues deemed essential for CM boundaries based on interactions observed between the HORMA domain and CMs in structural models generated with Alpha Fold 2.3. Protein models predicted with Alpha Fold 2.3 were analyzed with PyMOL and ChimeraX (Meng *et al*. 2023) in regards of conserved structures and residues in the interaction site between ecHOP1 and ecRED1 and their truncations. 18 models were chosen for further evaluation of the interaction between ecHOP1 HORMA domain and the putative ecRED1 CMs. Models were selected according to their rank and confidence of the model, which is indicated by the PAE-scores. The best ranking models generally have higher confidences and less clashes between amino acids, hence these were selected preferentially. Due to the varying numbers of models that show interactions between the HORMA domain and the putative ecRED1 CMs, the number of evaluated models per CM differs. Interactions between ecHOP1 and EcRed1 were determined by visualization of hydrogen bonds and polar contacts in a radius of 4 Å around the CM-residues to the HORMA domain. Furthermore, presence of hydrophobic interactions, π -stacking, or other interactions like π – Met –, π – Thiol, and π – Cation/Anion were investigated. Predicted H-bonds and salt bridges were further confirmed by PDBePISA(Krissinel Henrick 2007), which predicts interactions between two chains of a model computationally.

### Cloning of Expression Constructs

Sequences of full-length *Ectocarpus* ecHOP1 and ecRED1 were codon optimized for E. coli recombinant protein expression and ordered as gBlocks Gene Fragments (GenScript). All desired full-length and truncations were amplified by Phusion Flash High-Fidelity PCR using gene-specific primers (Supplementary Table 2; IDT DNA), and subsequent assembling via Gibson Assembly Cloning within the desired pCOLI- expression vectors (Altmannova *et al*. 2021).

### Protein expression and Purification

See Supplementary Methods

### SEC-MALS analysis

Triplicates at 1 mg/mL (approximately 50 µM) were loaded onto a Superrose 6 5/150 analytical size exclusion column (Cytiva) equilibrated in SEC-MALS buffer (50 mM HEPES, pH 7.5, 1 mM TCEP, 150 mM NaCl) attached to a 1260 Infinity II LC System (Agilent). MALS was completed with a Wyatt DAWN detector attached in line with the size exclusion column. Calibration and normalization was completed with monomeric BSA at 1 mg/mL in SEC-MALS buffer.

### Pull-Down Assays

Strep-TactinXT pull-down assays were performed with purified EcHOP1 HORMA domain with a Twin-Strep-tag® and His6-MBP tagged EcRED1 putative CM peptides. 250 µL reactions with 20 µg of each bait and prey protein in wash buffer (100 mM Tris pH 8.0, 300 mM NaCl, 1mM EDTA, 1mM TCEP) were incubated with 5 µL of magnetic Strep-TactinXT beads (IBA Lifesciences) for 20 minutes at 4°C with periodic gentle mixing. Beads were then washed two times with 500 µL of wash buffer. Bound proteins were eluted by resuspending the beads in 125 µL of elution buffer (100 mM Tris pH 8.0, 300 mM NaCl, 1mM EDTA, 1mM TCEP, 5% glycerol, 50 mM biotin) and subsequent incubation for 15 min at 4°C with gentle mixing throughout, repeating this process three times. To analyze the remaining proteins bound to the beads, a denaturing elution was performed by resuspending the beads in 30 µL of SDS loading dye and boiling for five minutes at 95°C. Samples for SDS-PAGE were prepared at a 2:1 ratio of sample and SDS loading dye, then boiled for five minutes at 95 °C. The gels were visualized with Coomassie staining (Der Blaue Jonas, Biozol), and Western Blot, using a 1:10000 dilution of anti MBP-tag antibody in 5% milk in 1x TBST, pH 7.6 (New England BioLabs), washed in 1x TBST, pH 7.6, followed by secondary anti-mouse antibody at 1:20000 for 20 minutes RT in 3% BSA in 1x TBST, pH 7.6 (Merck).

### Isothermal Titration Calorimetry (ITC)

ITC experiments were completed with a MicroCal PEAQ-ITC (Malvern Panalytical). All protein samples were extensively dialyzed in ITC buffer (50 mM HEPES pH 7.5, 150 mM NaCl, 1 mM EDTA) in Slide-A-Lyzer™ Dialysis Devices, 3.5K MWCO (ThermoFisher) at 4 °C O/N. Final concentrations were validated for both HORMA and all CMs, at 25 µM and 350 µM respectively. A ratio of 1:40 was used due to the predicted low binding affinity between the HORMA domain and CMs, as also recommended by the equipment manufacturer. The titrants (CMs, individually) were injected into the titrand (HORMA) at a starting injection of 0.4 µL, followed by 12x 3 µL injection volumes, with 180 second spacing, and spinning at 500 rpm, while maintaining a temperature of 25 °C. Control runs were completed with the same parameters, whether the titrant or titrand was measured in the addition of the ITC buffer. Buffer runs were also extensively evaluated to ensure minimal non-specific heat signals would not be acquired. All experiments were completed in triplicate to ensure reproducibility.

### Electrophoretic Mobility Shift Assays

Triplicate EMSAs were completed at a constant nucleic acid (167 bp Widom sample, dsDNA, ssDNA) concentration of 50 nM and seven samples of each protein ranging from 25 nM to 2 µM. Mononucleosomes were prepared as previously described (Rousova *et al*. 2021). The samples were incubated for approximately 30 minutes at 4 °C, then loaded onto a 0.8% agarose gel in 0.2% TBE Buffer. The gel was run for 2 h at 4 °C, not exceeding a power of 60 V, then post-stained with SYBRGold (Invitrogen). Gels were imaged using a ChemiDocMP (Bio-Rad Inc). Nucleic acid depletion in each lane was quantitated with the imager software, using measurements of triplicate of nucleic acid alone for each individual gel as a baseline. Binding curves were fitted using Prism software and the following algorithm (Y = Bmax*Xh/(KDh + Xh)).

## Acknowledgements

We would like to thank Remy Luthringer for supplying the various *Ectocarpus* tissue for these studies and for providing microscopic images of the various life stages of *Ectocarpus*. We also would like to thank Fabian Haas for their support with RNAseq analysis and assistance in generating boxplots, and Josué Barrera-Redondo for their contributions to the initial identification of ecHOP1 and ecRED1 in the *Ectocarpus* genome. We thank Veronika Altmannova for help with running and analysing the EMSAs. Many thanks to Jennifer Jüngling, Rahmiye Kürkcü and Susanne Astrinidis for outstanding technical support in the Weir laboratory. We thank Gerben Vader (Princess Maxima Cancer Centre, Utrecht) for comments on the manuscript. JRW is supported by the Max Planck Society and the German Research Foundation (Grant WE 6513/2-1). SMC is supported by the Max Planck Society, The European Research Council (Grant 864038), The Moore Foundation (Grant 11489) and the Bettencourt Foundation.

## CRediT Author Statement

EIK - Conceptualization, Methodology, Validation, Formal analysis, Investigation, Visualization, Supervision, Writing - Original Draft, Writing - Review & Editing LST - Methodology, Validation, Formal analysis, Investigation, Visualization LE - Methodology SMC - Conceptualization, Supervision, Project administration, Funding acquisition, Writing - Review & Editing JRW - Conceptualization, Formal analysis, Supervision, Visualization, Project administration, Funding acquisition, Writing - Review & Editing

## Supplementary Data

**Supplementary Table 1.**
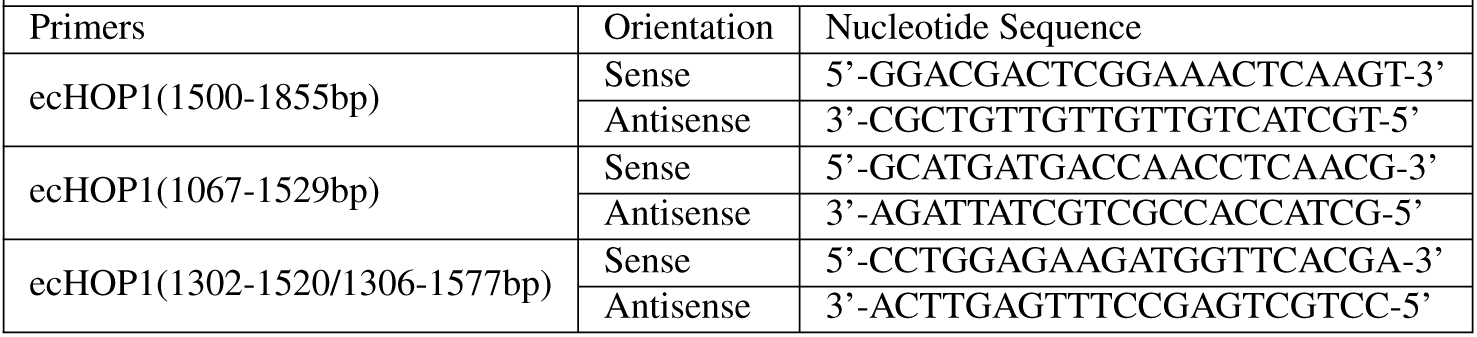
Oligonucleotide primers for ecHOP1 RT-PCR.

**Supplementary Table 2.**
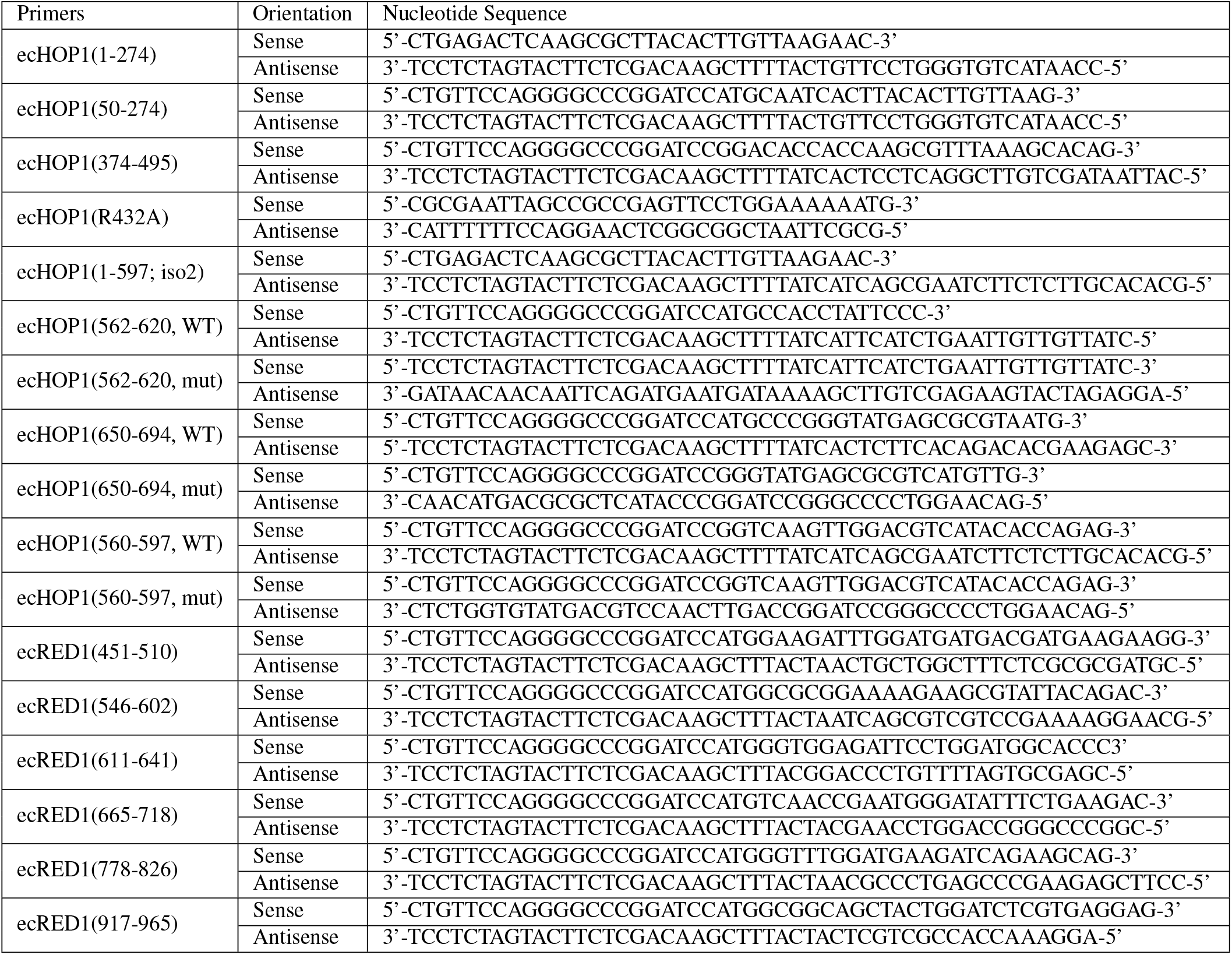
Oligonucleotide primers for ecHOP1 and ecRED1 Cloning.

**Supplementary Table 3.** Expression Vectors for Recombinant Protein Expression.

|  |  |  |  |
| --- | --- | --- | --- |
| Plasmid ID | Description | Domain Boundaries / Key Features | Reference |
| pWL1566 | StrepII-ecHOP1 1-694 | FL; isoform 1 | this study |
| pWL2556 | His6SUMO-ecHOP1 1-597 | FL; isoform 2 | this study |
| pWL1776 | StrepII-ecHOP1 1-274 | HORMA (WT) with StrepII tag | this study |
| pWL2555 | His6SUMO-ecHOP1 1-274 | HORMA (WT) with His6SUMO tag | this study |
| pWL2559 | His6SUMO-ecHOP1 50-274 | HORMA (dN-helix) | this study |
| pWL2562 | His6SUMO-ecHOP1 375-462; WT | wHTH (WT) | this study |
| pWL2566 | His6SUMO-ecHOP1 375-462; R432A | wHTH (R432A) | this study |
| pWL2546 | His6MBP-ecHOP1 562-620; WT | ecHOP1 CM1-1 WT | this study |
| pWL2547 | His6MBP-ecHOP1 562-620; R582-583A, K591A, R592A | ecHOP1 CM1-1 mutant | this study |
| pWL2550 | His6MBP-ecHOP1 650-694; WT | ecHOP1 CM2-1 WT | this study |
| pWL2551 | His6MBP-ecHOP1 650-694; R671-672A, K687-688A | ecHOP1 CM2-1 mutant | this study |
| pWL2552 | His6MBP-ecHOP1 560-597; WT | ecHOP1 CM1-2 WT | this study |
| pWL2553 | His6MBP-ecHOP1 560-597; R590-591A | ecHOP1 CM1-2 mutant | this study |

**Supplementary Figure 1.**
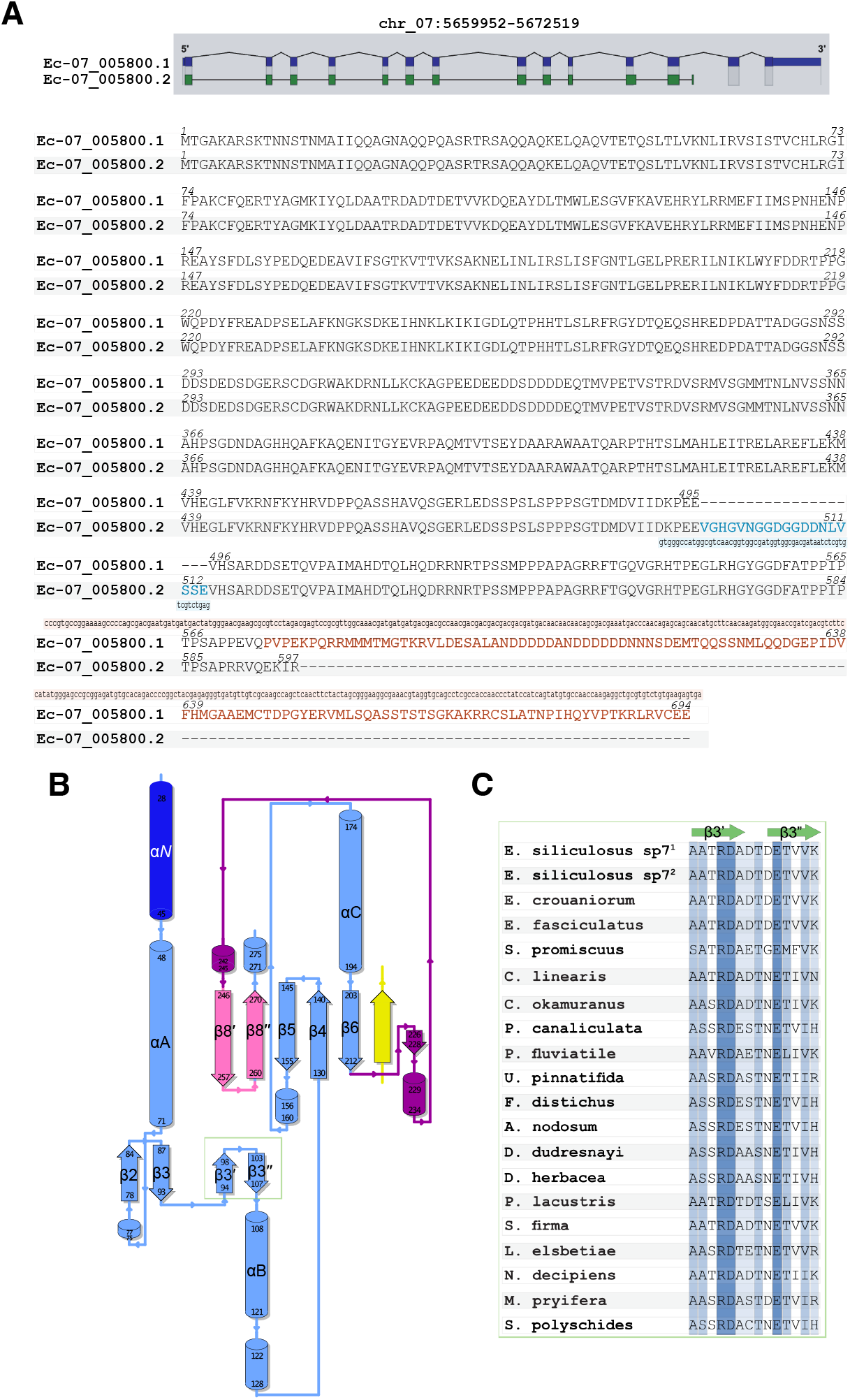
A) MSA ecHOP1 transcriptional isoforms. (top) Genome annotation figure of thw two ecHOP1 isoforms from ORCAE. (bottom) Alignment between the two transcriptional isoforms of ecHOP1 (Ec-07_005800.1 for isoform #1 and Ec-07_005800.2 for isoform #2) show high conservation between the two. Interestingly, there is an insertion between a.a. 495-514 in isoform 2 (blue), as well as a C-terminal extension following a.a. 574 in isoform 1 (orange). Although the insertion in isoform 2 does not computationally demonstrate to influence the protein structure, the extension in isoform 1 contains the two putative CMs. The DNA sequences for each isoform were acquired through ORCAE, and the translated sequences were utilized for MSA. The MSA was generated with JalView and aligned with MAFFT using default settings. B) Topology map of ecHOP1 HORMA domain, coloured as in Figure 2. C) MSA of the brown algal HOP1 orthologs, highlighting the novel β3 region (residues 94-107). A color gradient from light-to-dark was implemented to indicate percentage of conservation. The alignment was generated using MAFFT and curated in Jalview.

**Supplementary Figure 2.**
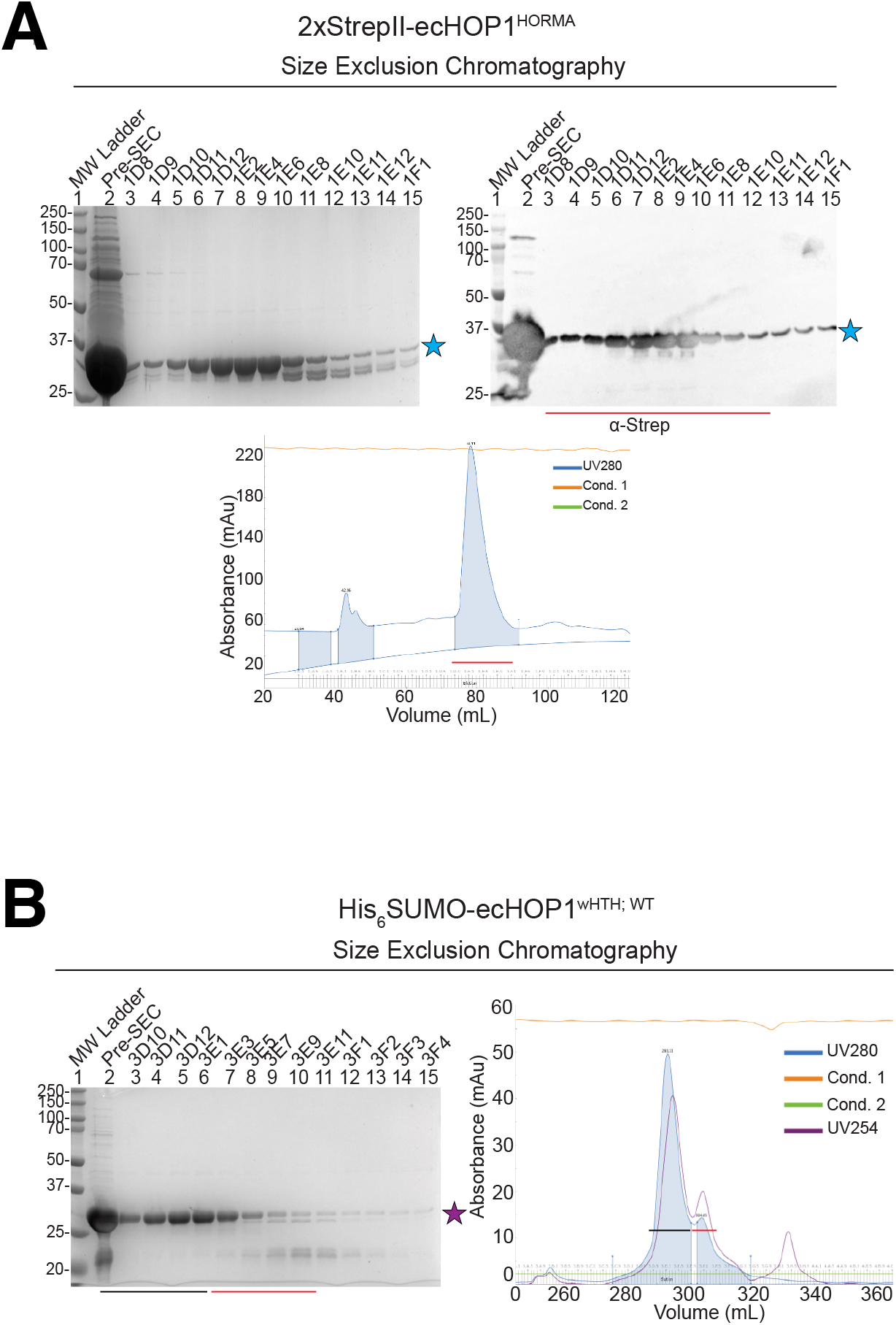
Purification of ecHOP1 proteins. (A) Purification of 2xStrepII-ecHOP1HORMA B) Purification of His6SUMO-ecHOP1wHTH.

**Supplementary Figure 3.**
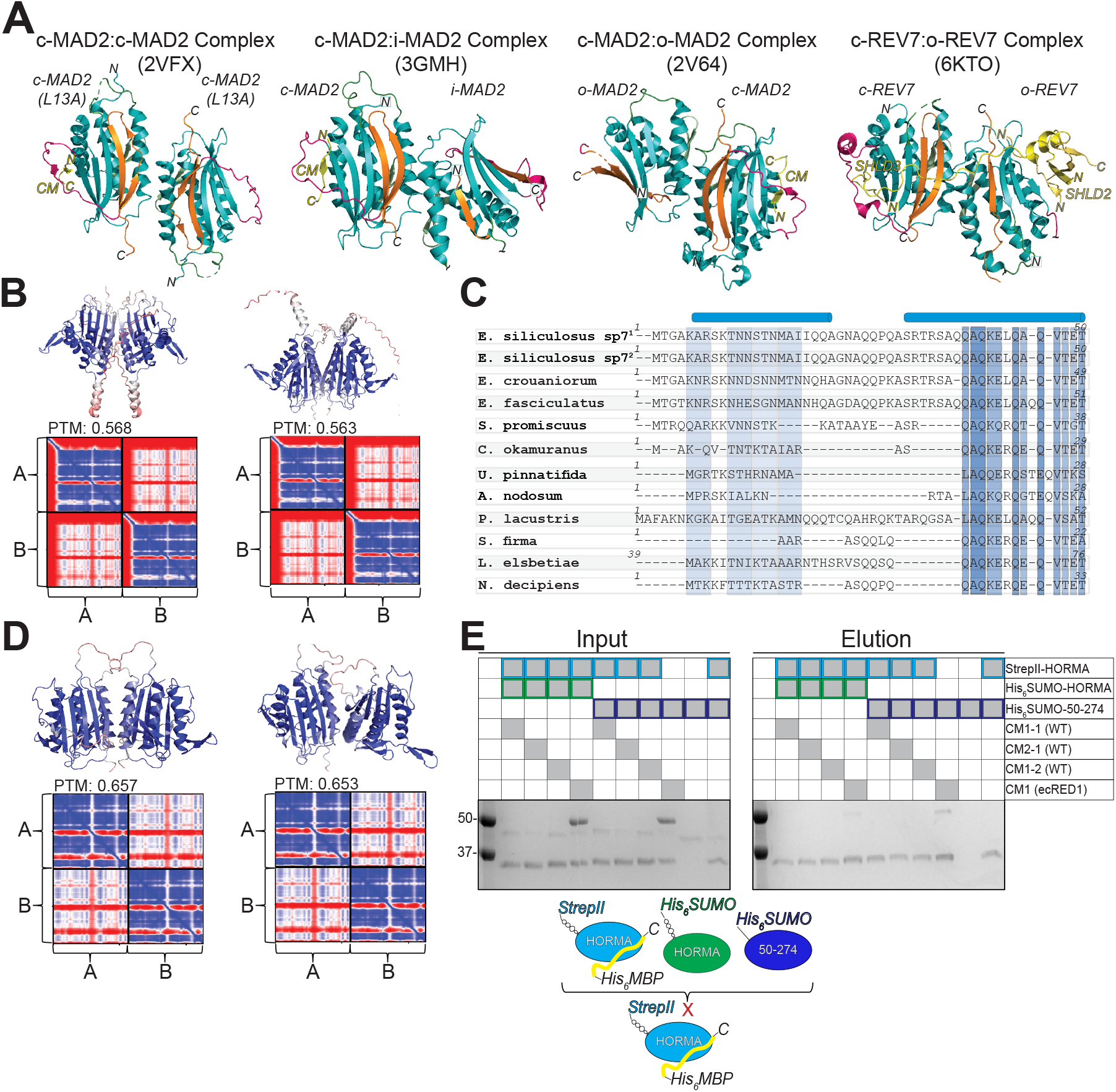
Homodimerization of ecHOP1-HORMA. (A) Experimentally determined structures of the c-MAD2L13A:c-MAD2L13A, (PDB 2VFX), c-MAD2:i-MAD2 (PDB 3GMH), c-MAD2:o-MAD2 (PDB 2V64), and c-REV7:o-REV7 complex (PDB 6TKO).These complexes involve one closed MAD2 stabilized by either an open MAD2 or an unbound CM from i-MAD2. REV7 dimerization, critical for SHLD2 binding, relies on conserved residues, including the key Arg124 in HsREV7. (B) AF2 models of ecHOP1 (a.a. 1–274) folding prior to the HORMA domain. However, steric clashes suggest that HORMA domain dimerization in ecHOP1 is unlikely. (C) MSAs of the N-terminal α-helix across brown algae, excluding species with incomplete annotations. (D) Models omitting the N-terminal α-helix show improved structural confidence and no steric clashes, supporting the hypothesis that the α-helix inhibits HORMA domain dimerization. (E) Pull-down assays with a closed 2xStrepII-ecHOP1HORMA prey (incubated with known CM interactors) and His6SUMO-HORMA or His6SUMO-ecHOP150-274 baits failed to validate ecHOP1 HORMA dimerization. Full-length protein may be necessary to form a self-closed HORMA with one CM and bind another HORMA via the second CM.

**Supplementary Figure 4.**
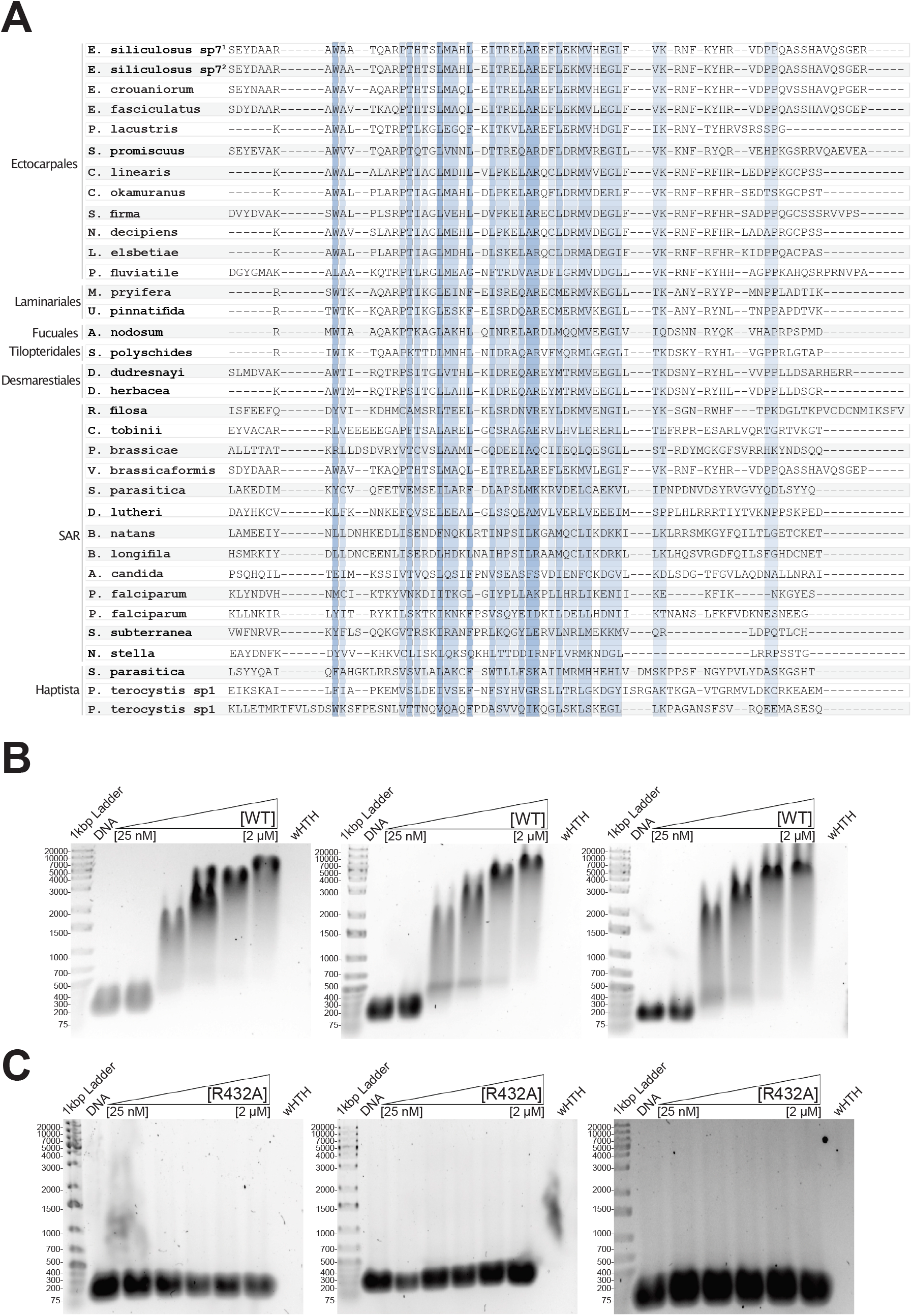
ecHOP1 wHTH domain and DNA binding properties. (A) MSA of HOP1 wHTH within Stramenopiles. B) EMSAs of WT ecHOP1 wHTH on 167 bp dsDNA B) EMSAs of R432A mutant ecHOP1 wHTH on 167 bp dsDNA.

**Supplementary Figure 5.**
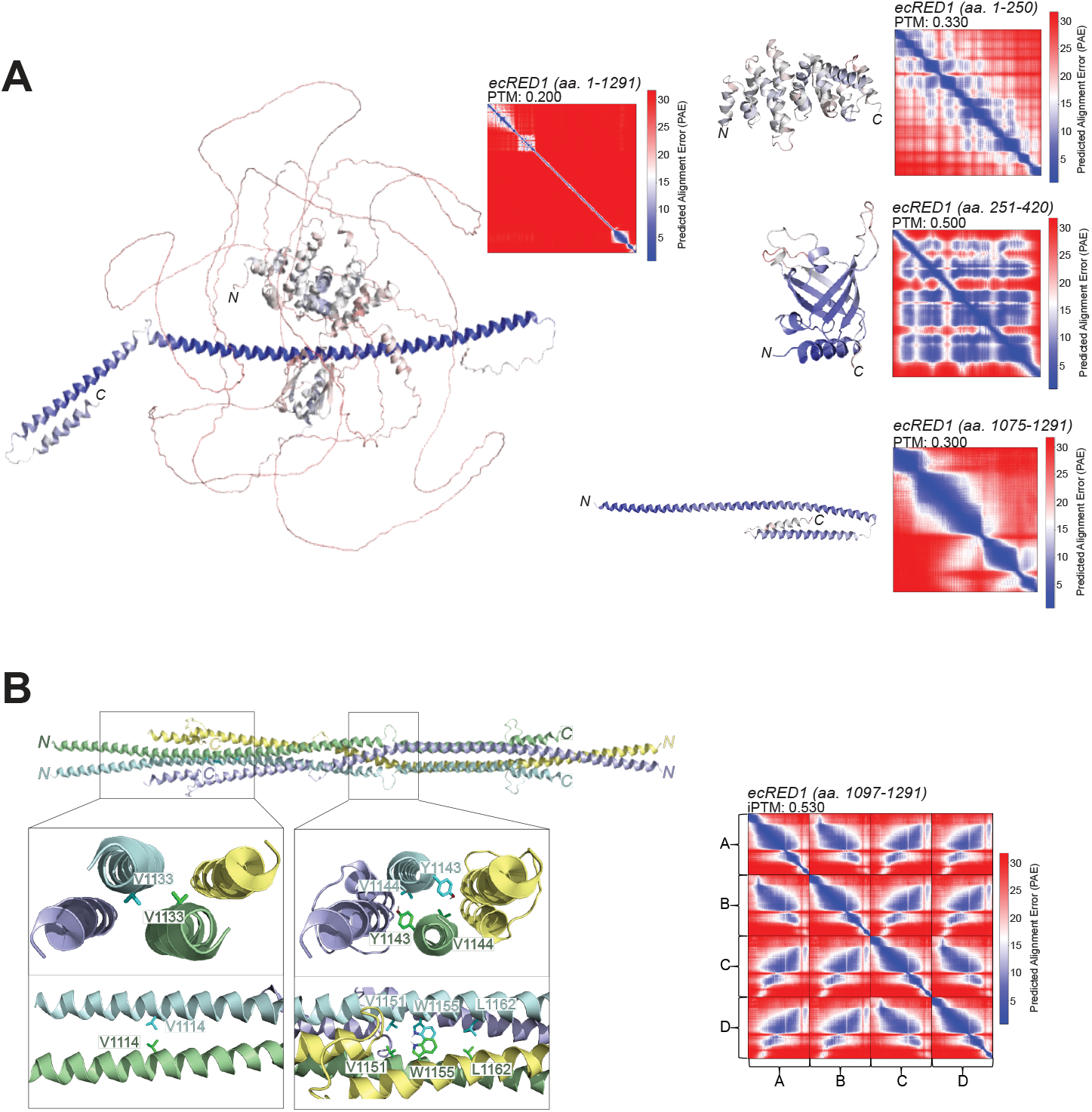
ecRED1 structure modelling and validation. (A)AlphaFold2 model of ecRED1 domains with PAE plots. B) AlphaFold2 models of ecRED1 coiled-coil domain as a tetramer.

**Supplementary Figure 6.**
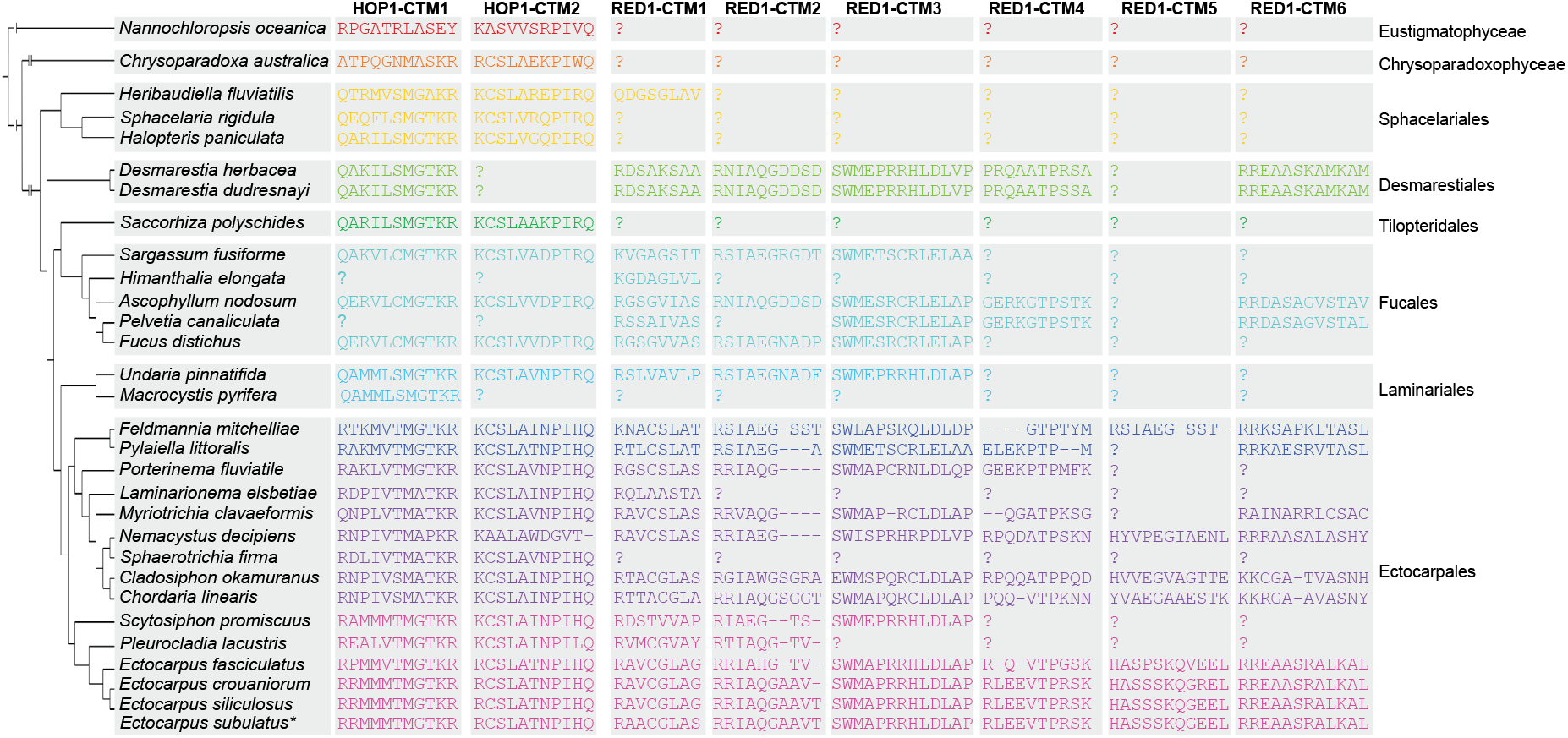
Closure Motif Phylogeny MSA and phylogenetic analysis of brown algal HOP1 and RED1 orthologs reveal a lineage-specific rise of CMs. Absence of specific CMs in certain organisms is due to poor genome annotations or lack of conservation in relevant regions. In Ectocarpales, ecHOP1 and ecRED1 CMs are highly conserved, with additional ecRED1 CMs emerging in Desmarestiales. HOP1 orthologs in Eustigmatophyceaes and Chrysoparadoxophyceaes show two well-conserved CMs, suggesting an early addition of CMs for regulatory functions. The alignment was generated using MAFFT and curated in Jalview to identify conserved K/R residues defining CMs.

## Supplementary Methods

### Overexpression and purification of ecHOP1 HORMA domain

ecHOP1HORMA was expressed as a 3C HRV cleavable N-terminal StrepII-tag fusion in BL21(DE3) competent cells. The cultures were grown at 37 °C until an OD600 of 0.6 was achieved. Protein expression was induced with 500 µM IPTG at 16 °C for approximately 16 hours. The cells were centrifuged at 15,000 rpm for 10 min, and the pellets were resuspended in cell lysis buffer (100 mM KH2PO4 pH 7.3, 250 mM NaCl, 1 mM MgCl2, 10 mM arginine, 10 mM glutamate, 10 mM imidazole, 0.1% Triton-X 100). The cells were then lysed using an EmulsiFlex C3 (Avestin) in the presence of SERVA protease (25 µg/mL), AEBSF protease (25 µg/mL), and DNAseI (10 µg/mL) (ThermoFisher) and cleared through ultracentrifugation at 40,000 rpm for one hour at 4 °C (Beckman Coulter). The supernatant was then passed through a 5 mL StrepTactinXT 4 flow column (IBA Lifesciences), previously equilibrated in wash buffer (100 mM KH2PO4 pH 7.3, 250 mM NaCl, 1 mM EDTA, 1 mM MgCl2, 1 mM TCEP, 10 mM arginine, 10 mM glutamate). The column was washed with an ATP wash buffer (100 mM KH2PO4 pH 7.3, 250 mM NaCl, 1 mM EDTA, 1 mM MgCl2, 1 mM TCEP, 10 mM arginine, 10 mM glutamate, 1 µM ATP), followed by a high salt wash buffer (100 mM KH2PO4 pH 7.3, 800 mM NaCl, 1 mM EDTA, 1 Mm MgCl2, 1 mM TCEP, 10 mM arginine, 10 mM glutamate) to eliminate non-specific contaminants such as chaperones and DNA. The protein was then eluted with elution buffer (100 mM KH2PO4 pH 7.3, 800 mM NaCl, 1 mM EDTA, 1 mM MgCl2, 1 mM TCEP, 10 mM arginine, 10 mM glutamate, 50 mM biotin), concentrated on an Amicon concentrator (30 MWCO; ThermoFisher) and loaded a Superdex 200 16/600 (Cytiva) pre-equilibrated in wash buffer. Biochemically pure fractions were pooled, flash frozen in N2(l) and stored in -80 °C for future use. ecHOP1HORMA and ecHOP150-274 were also expressed as a 3C HRV cleavable N-terminal His6SUMO-tag fusion in BL21(DE3) competent cells. The cultures were grown at 37 °C until an OD600 of 0.6 was achieved. Protein expression was induced with 500 µM IPTG at 16 °C for approximately 16 hours. The cells were centrifuged at 15,000 rpm for 10 min, and the pellets were resuspended in cell lysis buffer (100 mM KH2PO4 pH 7.3, 250 mM NaCl, 1 mM MgCl2, 10 mM arginine, 10 mM glutamate, 10 mM imidazole, 0.1% Triton-X 100). The cells were then lysed using an EmulsiFlex C3 (Avestin) in the presence of SERVA protease (25 µg/mL), AEBSF protease (25 µg/mL), and DNAseI (10 µg/mL) (ThermoFisher) and cleared through ultracentrifugation at 40,000 rpm for one hour at 4 °C (Beckman Coulter). The supernatant was then passed through a 5 mL HisTrap column (Cytiva), previously equilibrated in wash buffer (100 mM KH2PO4 pH 7.3, 250 mM NaCl, 1 mM EDTA, 1 mM MgCl2, 10 mM arginine, 10 mM glutamate, 10 mM imidazole). The column was extensively washed in wash buffer, then the protein was then eluted with elution buffer (100 mM KH2PO4 pH 7.3, 800 mM NaCl, 1 mM EDTA, 1 mM MgCl2, 10 mM arginine, 10 mM glutamate, 250 mM imidazole), and buffer exchanged via dialysis (3.5 MWCO; ThermoFisher) in low salt buffer (100 mM KH2PO4 pH 7.3, 50 mM NaCl, 1 mM EDTA, 1 mM MgCl2, 1 mM TCEP, 10 mM arginine, 10 mM glutamate). The protein was then loaded onto a HiTrap S column (Cytiva) pre-equilibrated in low salt buffer. A stepwise gradient with high salt buffer (100 mM KH2PO4 pH 7.3, 1 M NaCl, 1 mM EDTA, 1 mM MgCl2, 1 mM TCEP, 10 mM arginine, 10 mM glutamate) was used to elute the protein and non-specific contaminants from the column. The flow through, wash, and elution gradients up to 25% were pooled, concentrated on an Amicon concentrator (10 MWCO; ThermoFisher) flash frozen in N2(l) and stored in -80 °C for future use.

### Overexpression and purification of ecHOP1 wHTH domain

ecHOP1wHTH was expressed as a 3C HRV cleavable N-terminal His6SUMO-tag fusion in BL21(DE3) competent cells. The cultures were grown at 37 °C until an OD600 of 0.6 was achieved. Protein expression was induced with 500 µM IPTG at 16 °C for approximately 16 hours. The cells were centrifuged at 15,000 rpm for 10 min, and the pellets were resuspended in cell lysis buffer (100 mM KH2PO4 pH 7.3, 250 mM NaCl, 1 mM MgCl2, 10 mM arginine, 10 mM glutamate, 10 mM imidazole, 0.1% Triton-X 100). The cells were then lysed using an EmulsiFlex C3 (Avestin) in the presence of SERVA protease (25 µg/mL), AEBSF protease (25 µg/mL), and DNAseI (10 µg/mL) (ThermoFisher) and cleared through ultracentrifugation at 40,000 rpm for one hour at 4 °C (Beckman Coulter). The supernatant was then passed through a 5 mL HisTrap column (Cytiva), previously equilibrated in wash buffer (100 mM KH2PO4 pH 7.3, 250 mM NaCl, 1 mM EDTA, 1 mM MgCl2, 10 mM arginine, 10 mM glutamate, 10 mM imidazole). The column was extensively washed in wash buffer, then the protein was then eluted with elution buffer (100 mM KH2PO4 pH 7.3, 800 mM NaCl, 1 mM EDTA, 1 mM MgCl2, 10 mM arginine, 10 mM glutamate, 250 mM imidazole), and buffer exchanged via dialysis (3.5 MWCO; ThermoFisher) in low salt buffer (100 mM KH2PO4 pH 7.3, 50 mM NaCl, 1 mM EDTA, 1 mM MgCl2, 1 mM TCEP, 10 mM arginine, 10 mM glutamate). The protein was then loaded onto a HiTrap S column (Cytiva) pre-equilibrated in low salt buffer. A stepwise gradient with high salt buffer (100 mM KH2PO4 pH 7.3, 1 M NaCl, 1 mM EDTA, 1 mM MgCl2, 1 mM TCEP, 10 mM arginine, 10 mM glutamate) was used to elute the protein and non-specific contaminants from the column. The flow through, wash, and elution gradients up to 25% were pooled, concentrated on an Amicon concentrator (10 MWCO; ThermoFisher) flash frozen in N2(l) and stored in -80 °C for future use.

### Overexpression and purification of ecRED1 CMs

All ecRED1CMs were expressed as a 3C HRV cleavable N-terminal His6MBP-tag fusion in BL21(DE3) Artic Express competent cells. The cultures were grown at 30 °C until an OD600 of 0.6 was achieved. Protein expression was induced with 500 µM IPTG at 12 °C for approximately 16 hours. The cells were centrifuged at 15,000 rpm for 10 min, and the pellets were resuspended in cell lysis buffer (50 mM HEPES pH 7.5, 300 mM NaCl, 1 mM TCEP, 0.1% Triton-X 100). The cells were then lysed using an EmulsiFlex C3 (Avestin) in the presence of SERVA protease (25 µg/mL), AEBSF protease (25 µg/mL), and DNAseI (10 µg/mL) (ThermoFisher) and cleared through ultracentrifugation at 40,000 rpm for one hour at 4 °C (Beckman Coulter). The supernatant was then passed through a 5 mL HisTrap column (Cytiva), previously equilibrated in wash buffer (50 mM HEPES pH 7.5, 300 mM NaCl, 1 mM EDTA, 1 mM TCEP). The column was extensively washed in wash buffer, then the protein was then eluted with elution buffer (100 mM KH2PO4 pH 7.3, 800 mM NaCl, 1 mM EDTA, 1 mM MgCl2, 10 mM arginine, 10 mM glutamate, 250 mM imidazole), and concentrated on an Amicon concentrator (10 MWCO; ThermoFisher) and loaded a Superdex 200 16/600 (Cytiva) pre-equilibrated in wash buffer. Biochemically pure fractions were pooled, flash frozen in N2(l) and stored in -80 °C for future use.

## Notes

### Competing Interest Statement

The authors have declared no competing interest.

